# Regulatory kinase genetic interaction profiles differ between environmental conditions and cellular states

**DOI:** 10.1101/735720

**Authors:** Siyu Sun, Anastasia Baryshnikova, Nathan Brandt, David Gresham

## Abstract

Cell growth and quiescence in eukaryotic cells is controlled by an evolutionarily conserved network of signaling pathways. Signal transduction networks operate to modulate a wide range of cellular processes and physiological properties when cells exit proliferative growth and initiate a quiescent state. How signaling networks function to respond to diverse signals that result in cell cycle exit and establishment of a quiescent state is poorly understood. Here, we studied the function of signaling pathways in quiescent cells using global genetic interaction mapping in the model eukaryotic cell, *Saccharomyces cerevisiae* (budding yeast). We performed pooled analysis of genotypes using molecular barcode sequencing to test the role of ∼3,900 gene deletion mutants and ∼11,700 pairwise interactions between all non-essential genes and the protein kinases *TOR1, RIM15, PHO85* in three different nutrient-restricted conditions in both proliferative and quiescent cells. We detect nearly five-fold more genetic interactions in quiescent cells compared to proliferative cells. We find that both individual gene effects and genetic interaction profiles vary depending on the specific pro-quiescence signal. The master regulator of quiescence, *RIM15* shows distinct genetic interaction profiles in response to different starvation signals. However, vacuole-related functions show consistent genetic interactions with *RIM15* in response to different starvation signals suggesting that *RIM15* integrates diverse signals to maintain protein homeostasis in quiescent cells. Our study expands genome-wide genetic interaction profiling to additional conditions, and phenotypes, highlighting the conditional dependence of epistasis.

## Introduction

Most cells spend the majority of their lifetime in a quiescent state defined as the temporary and reversible absence of proliferation (O’Farrell 2011; Lemons et al. 2010; Valcourt et al. 2012). Quiescence requires exit from the mitotic cell division cycle and initiation of a distinct G0 cell cycle phase, during which cells remain viable and maintain the capacity to re-initiate proliferative growth (Valcourt et al. 2012). In multicellular organisms development, tissue renewal and long term survival is dependent upon the persistence of stem cells that are quiescent, but retain the ability to re-enter the cell cycle to self-renew, or to produce progeny that can differentiate and re-populate the tissue (Miles and Breeden 2017). Exit from quiescence, and initiation of aberrant proliferation, is a hallmark of cancer (Hanahan and Weinberg 2011; Miles and Breeden 2017). Conversely, many cancer-related deaths are the result of quiescent tumor cells that are resistant to therapeutics and underlie tumor recurrence (Yano et al. 2017; Borst 2012). Thus, understanding cellular quiescence and how cells regulate the transition between proliferative and quiescent states is of fundamental importance to our understanding of cellular homeostasis and disease.

Cells exit the cell cycle and enter quiescence when they are deprived of essential nutrients or growth factors (Valcourt et al. 2012; Daignan-Fornier and Sagot 2011; Klosinska et al. 2011). The quiescent state of the model eukaryotic organism, *Saccharomyces cerevisiae* (budding yeast) shares many important features with that of higher organisms (Valcourt et al. 2012; Dhawan and Laxman 2015), including cell cycle arrest, condensed chromosomes, reduced rRNA synthesis and protein translation, and increased resistance to stress. Therefore, the mechanisms that regulate cell cycle arrest and the establishment, maintenance and exit from a quiescent state, as well as the physiological processes associated with this state, are likely to be shared across eukaryotic cells.

Studies of quiescence in yeast typically examine stationary-phase cells, namely cells grown to saturation in rich, glucose-containing medium (Gray et al. 2004; Young et al. 2017). Such cells first exhaust available glucose through fermentative metabolism and then, following the diauxic shift, switch to respiration using ethanol as carbon source. Upon exhaustion of ethanol cells enter quiescence. However, in addition to carbon starvation, yeast cells can respond to a variety of nutrient starvations by exiting the cell cycle and initiating quiescence (Lillie and Pringle 1980; Gresham et al. 2011; Klosinska et al. 2011). Starvation for essential nutrients including nitrogen, phosphorus and sulfur result in many of the same characteristics as carbon starved cells including arrest as unbudded cells, thickened cell walls, increased stress resistance and an accumulation of storage carbohydrates (Lillie and Pringle 1980; Schulze et al. 1996). Although under laboratory conditions yeast primarily experience carbon starvation, in the wild yeast are likely to experience a diversity of nutrient deprivations. How the cell integrates these diverse signals to mount the same physiological response, and establish cellular quiescence, remains poorly understood.

The ability of stationary phase yeast cells to maintain viability has also been used as a model for the chronological aging. Chronological lifespan (CLS) has been defined as the time a yeast cell can survive in a non-dividing, quiescent-like state (Fabrizio and Longo 2003; Kaeberlein 2010; Walter, Matter, and Fahrenkrog 2014). Therefore, CLS is closely related to the proportion of quiescent cells in stationary phase cultures because non-quiescent cells have a reduced ability to reenter the cell cycle (Allen et al. 2006; Walter, Matter, and Fahrenkrog 2014). Cells with a shortened CLS have reduced reproductive capacity upon replenishment of nutrients, while cells with an increased CLS have enhanced reproductive capacity under the same condition (Garay et al. 2014). Identification of genes that mediate CLS in yeast under different nutrient restrictions is potentially informative about the regulation of aging in higher organisms.

Genotype has a profound impact on the regulation of quiescence. Many studies of survival in stationary-phase cells, and their application to the study of CLS, have been conducted using auxotrophic strains. However, starvation for an engineered auxotrophic requirement results in a failure to effectively initiate a quiescent state and therefore leads to a rapid loss of viability (Boer et al. 2008; Gresham et al. 2011). This is likely due to the fact that yeast cells have not evolved a mechanism for sensing and responding to lab engineered auxotrophic requirements. Thus, the identification of mutants that suppress the rapid loss of viability upon undefined starvation in auxotrophic strains may be of limited relevance for understanding the regulation of quiescence and CLS. Previous studies of quiescence using prototrophic yeast cells, and defined starvations, have shown that the genetic requirements for quiescence differ depending on the nutrients for which the cell is starved (Gresham et al. 2011; Klosinska et al. 2011). However, whether the genes required for proliferation in nutrient-limited conditions are they the same set of genes that are required for programming quiescence is not known.

Multiple evolutionarily conserved nutrient sensing and signal transduction pathways, including the target of rapamycin complex I (TORC1), protein kinase A (PKA), AMP kinase (AMPK) and PHO85 pathways have been shown to regulate quiescence. The integrator of these diverse signalling pathways is thought to be the protein kinase RIM15, a great wall kinase with a mammalian homologue called microtubule associated serine/threonine like kinase (MASTL) (Castro and Lorca 2018). This regulator appears to be downstream of multiple signaling pathways and is required for the establishment of stationary phase. However, how different starvation signals are coordinately transduced via these pathways, and how RIM15 orchestrates the establishment of cellular quiescence is not known (de Virgilio 2012; Broach 2012).

The relationship between different cellular processes and pathways can be investigated using a variety of methods that identify physical and functional interactions. One efficient approach to defining interactions between genes and pathways is through genetic interaction mapping. A genetic interaction is a relationship between two genes in which the phenotype of the double mutant diverges from that expected on the basis of the phenotype of each single mutant (Costanzo et al. 2010; Tong et al. 2004). A genetic interaction can be informative of the functional relationship between the encoded products. Positive genetic interactions may be indicative of genes that exist within pathways or complexes whereas negative genetic interactions often reflect genes that function in parallel pathways or processes that converge on the same function. Extension of genetic interaction mapping to testing genome-wide interactions between defined alleles enables definition of a genetic interaction profile, comprising the set of negative and positive genetic interactions for a given gene. The systematic application of this approach has demonstrated that genes that share similar functions, or operate in the same pathway, usually share similar genetic interaction profiles. As such, the similarity of a genetic interaction profile between two genes (typically quantified as a correlation coefficient) is informative about the similarity between the two genes’ functions. The culmination of genome-wide genetic interaction mapping in budding yeast is a global genetic interaction similarity network that serves as a functionally informative reference map (Costanzo et al. 2010; Costanzo et al. 2016). The completion of this comprehensive genetic interaction map leads to two related questions: 1) to what extent are genetic interactions dependent on environmental conditions? and 2) can genome-wide genetic interaction mapping be expanded to other phenotypes? To date, genome-wide genetic interaction mapping in yeast has primarily been performed in a single condition and assayed using a single phenotype – colony growth in rich media. The extent to which genetic interactions defined by cell growth phenotypes depend on environmental conditions and the utility of using additional phenotypes in genetic mapping remains largely unexplored. A small number of studies suggest that functional relationships between genotype and phenotype are environmentally dependent (Díaz-Mejía et al. 2018; Jaffe et al. 2019; Bandyopadhyay et al. 2010) suggesting that a complete understanding of global genetic interaction networks requires identification of genetic interactions in multiple conditions and using multiple phenotypes.

Here, we have developed a robust method for quantifying phenotypes of pooled single and double mutants in different conditions using barcode sequencing. We applied this approach to quantify genetic requirements, and identify genetic interactions, in two different cellular states and three different nutritional conditions. Our experimental design entailed quantification of both fitness during proliferative growth and survival during prolonged defined starvation for each genotype. We find that the genetic requirements for quiescence differ depending on the nutrient starvation signal. Using genome-wide genetic interaction mapping for three key regulatory kinases, we find that these genes exhibit different interaction profiles in different growth conditions and in different cellular states. Finally, we find that the master regulator of quiescence, *RIM15* shows distinct genetic interaction profiles and regulates different functional groups in response to different starvation signals. However, vacuole-related functions show consistent negative genetic interactions with *RIM15* in response to different starvation signals suggesting that *RIM15* controls quiescence by integrating diverse signals to maintain protein homeostasis. Our study points to a rich spectrum of condition-specific genetic interactions that underlie cellular fitness and survival across a diversity of conditions and introduces a generalizable framework for extending genome-wide genetic interaction mapping to diverse conditions and phenotypes.

## Results

### Quantifying mutant fitness using pooled screens in diverse conditions

Cellular quiescence in yeast can be induced through a variety of nutrient deprivations, but whether establishment of a quiescent state in response to different starvation signals requires the same genetic factors and interactions is unknown. Therefore, we sought to test the specificity of gene functions and genetic interactions in quiescent cells induced in response to three natural nutrient starvations: carbon, nitrogen and phosphorus. The use of prototrophic yeast strains is essential for the study of quiescence as unnatural, or unknown, starvations can confound results and their interpretation (Boer et al, 2008; Gresham et al. 2011). Therefore, we constructed haploid prototrophic double mutant libraries using a modified synthetic genetic array (SGA) mating and selection method (**Fig EV1A**). Briefly, double mutant libraries were constructed using genetic crosses between the ∼4,800 non-essential gene deletion strains (Giaever et al. 2002) and query strains deleted for one of three genes encoding the catalytic subunit of different regulatory protein kinases: *TOR1, RIM15*, and *PHO85* (**methods**). In addition, we constructed a single mutant library using the same method by mating the gene deletion collection with a strain deleted for *HO*, which has no fitness defects in haploids. We confirmed the genotype and ploidy of the resulting three haploid double gene deletion libraries and the single mutant library using selective media and flow cytometry (**Fig EV1B**).

Previously, genome-wide genetic interaction mapping in yeast has been performed using colony growth phenotypes as a measurement of genotype fitness (Costanzo et al. 2010; Costanzo et al. 2016). In liquid cultures, the growth cycle of a population of microbial cells comprises a lag period, an exponential growth phase, and a subsequent period in which growth is no longer observed, known as stationary phase. Initiation of stationary phase is indicative of cell growth and cell cycle arrest due to starvation for an essential nutrient (de Virgilio 2012). To study each genotype over the complete growth cycle in liquid cultures, we first analyzed the four libraries (**Fig 1A**) in three different defined nutrient-restricted media: carbon-restriction (minimal media containing 4.4mM carbon), nitrogen-restriction (minimal media containing 0.8mM nitrogen), and phosphorus-restriction (minimal media containing 0.04 mM phosphorus) (**Table EV2, methods and materials**). The composition of these media ensures that following an exponential growth phase cells experience either carbon, nitrogen or phosphorus starvation, respectively. In each of the three conditions media, 1.5 × 10^8^ cells from each of the four libraries (**Fig 1A**) of pooled mutants was used to inoculate cultures (t = 0). In nitrogen- and phosphorus-restriction media, we observed that the starvation period commenced 24 hours after inoculation (**Fig EV1C**). Cells in carbon-restricted media underwent the diauxic shift after 24 hours, and reached stationary phase approximately 48 hours post inoculation (**Fig EV1C**). Beyond these time points we did not observe additional cell division or population expansion consistent with defined nutrient starvation and the initiation of quiescence.

**Figure 1.**
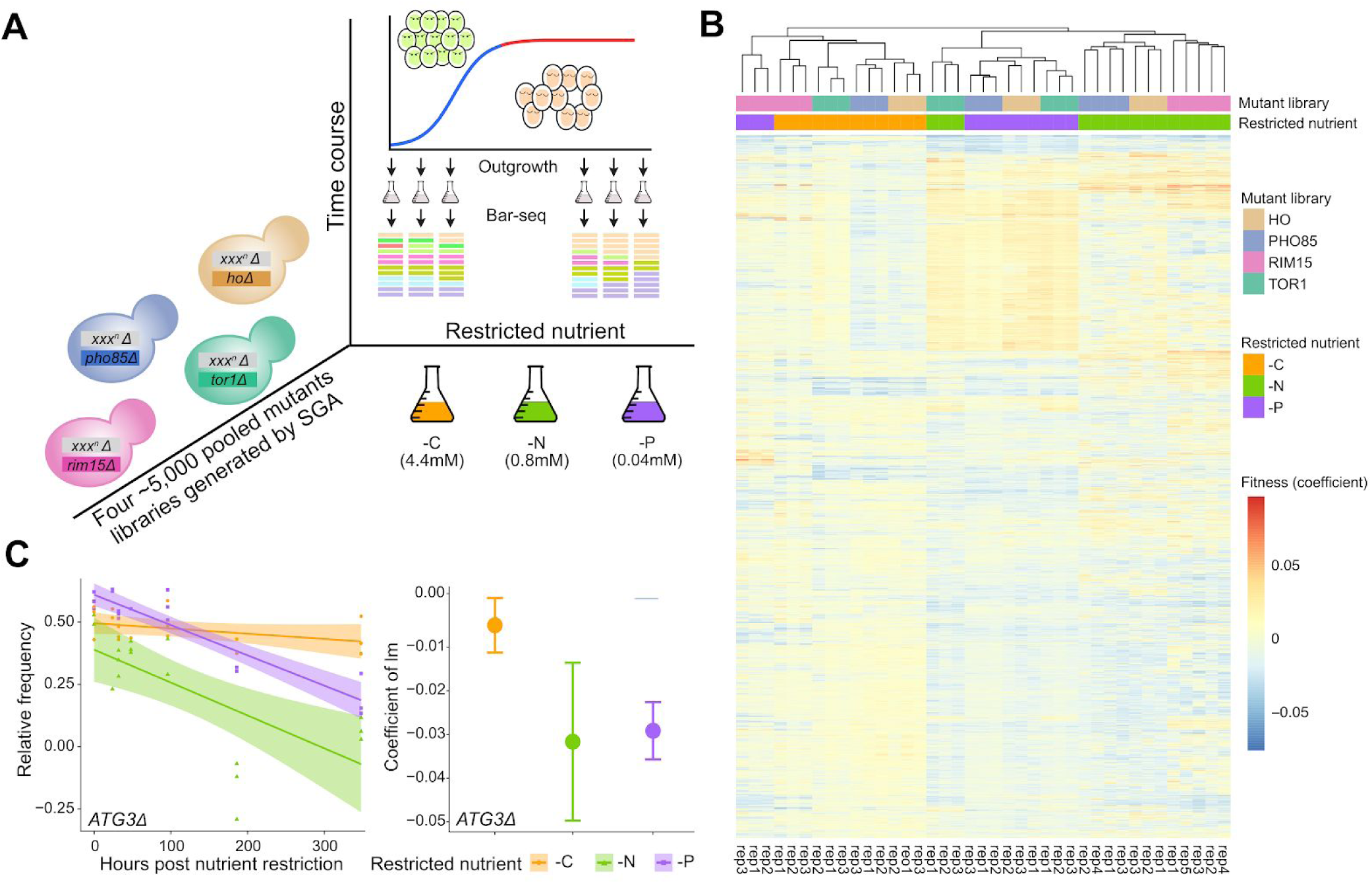
Robust fitness estimation over the entire population growth cycle using pooled mutant libraries and Bar-seq. **A)** Experimental design for multiplexed mutant survival assay using Bar-seq. The synthetic genetic array (SGA) method was used to construct four genome-wide double-mutant prototrophic libraries (**Fig EV1A**). The yeast deletion collection (*xxx*^*n*^*Δ::natMX*) was mated with query strains deleted for one of three genes that encode regulatory kinases important in quiescence: *TOR1(tor1Δ::kanMX), RIM15 (rim15Δ::kanMX)*, and *PHO85 (pho85Δ::kanMX)*. A control library was made by mating the deletion collection to a neutral gene deletion HO (*hoΔ::kanMX*) query mutant. To ensure library complexity, 1.5 × 10^8^ cells from each library was used to inoculate (t=0) cultures restricted for glucose (-C, 4.4mM), ammonium sulfate (-N, 0.8mM), and potassium phosphate (-P, 0.04mM) in 300mL cultures. The starvation period for -N and -P conditions commenced after 24 hours and after 48 hours for -C condition (**Fig EV1C**). At different time points we removed a ∼2 × 10^5^ cell sample from the culture and expanded the viable subpopulation using outgrowth in supplemented minimal media (**Table EV2**). DNA was isolated from the resulting outgrowth culture and the library composition was analyzed using Bar-seq. **B)** Representative gene (*ATG3*) for relative fitness estimation across the entire culturing period. The abundance of the *atg3Δ0* strain was determined at multiple timepoints on the basis of counts of its unique DNA barcode and fitness was determined using linear regression. Linear models (predicted value +/-95% CI) fit to the data are shown on the left, coloured by condition. The coefficient (slope) of each model is shown in the dot plot on the right, with a 95% confidence interval indicated as an error bar (**Materials and Methods**). **C)** Hierarchical clustering of mutant fitness profiles computed for each replicate separately across the entire culturing period. White indicates that the strain has not changed in fitness compared to wild-type, blue represents increased fitness and red represents decreased fitness. Culture conditions are indicated by color (orange: carbon limited; green: nitrogen limited; purple: phosphorus limited). Three kinases mutant libraries (*TOR1, RIM15, PHO85*) and one control library (*HO*) are shown.

To compare the fitness of each genotype over the complete growth cycle in each condition, a 1mL sample was removed from the culture at different time points and the subpopulation of viable cells expanded using 24-48 hours of outgrowth in supplemented minimal media (**Fig 1A, method and materials**). This step is required to enrich for mutants that survive proliferation and starvation and to deplete those that have undergone senescence. Using an identical outgrowth step at every time point, and determining the rate of change in the relative abundance of viable mutants in the outgrown population, accounts for growth rate differences between mutants during the outgrowth (Gresham et al. 2011). The abundance of each mutant in the heterogenous pool was estimated by sequencing DNA barcodes that uniquely mark each genotype using Bar-seq (Smith et al. 2009; Gresham et al., 2011; Robinson et al., 2014). In total, we studied the four libraries in the three conditions with between 3-5 independent experiments to account for biological and technical variability (total of 39 genetic screens).

To determine the fitness of each strain during the complete growth cycle, we initially applied linear regression modeling of the relative frequency of each mutant against time (t = 0, 24, 48, 96, 186, 368 hours) (**Fig EV1C**). To test the reproducibility of our fitness assay, we first estimated fitness for each biological replicate separately and used PCA analysis to identify poorly behaved libraries (**Appendix_Fig1**). Hierarchical clustering of the filtered libraries show that for all 39 experiments biological replicates cluster as nearest neighbors (**Fig 1B**). Different libraries cultured in the same medium tend to cluster together, indicating that environmental conditions are a major determinant of fitness effects (**Fig 1B**). In general, mutants in carbon-restricted media show the opposite fitness profile to that observed in nitrogen and phosphorus-restricted conditions. However, the *TOR1* library in nitrogen-restricted media and the *RIM15* library in phosphorus-restricted media were exceptions to this general trend (**Fig 1B**).

To quantify fitness, and the associated uncertainty (expressed as a 95% confidence interval) of each estimate, we performed model fitting for each library in each condition using all biological replicates. We identified numerous cases in which the fitness of a single mutant significantly differs between conditions. For example, deletion of the autophagy gene *ATG3* (*atg3Δ0 hoΔ0)*, results in reduced fitness in nitrogen- and phosphorus-restriction media, but not in carbon-restriction media (**Fig 1C**).

### Distinct cellular functions are required for quiescence in response to different nutrient starvation signals

The fitness of a genotype during proliferative growth in different media may differ from the survival of the genotype in response to a specific starvation signal. Previous genome-wide genetic analyses of quiescence quantified the survival of each mutant in the absence of specific essential nutrients but did not assess the effect of each gene deletion on cellular proliferation prior to starvation (Klosinska et al. 2011). We test whether the genetic requirements for proliferation in nutrient-restricted media and quiescence in response to starvation for the same nutrient are distinct we separately modeled the relative abundance of each genotype during the growth phase (i.e. from t = 0 to t = 24 hours) and during the starvation period (i.e. from t = 32 to t = 368 hours). This analysis distinguishes the effect of each gene deletion in two distinct physiological states: proliferation and quiescence. As cells do not generate progeny when starved we refer to the phenotype during the starvation phase as “survival” and phenotype during proliferation as “fitness” (**Fig 2A**). We find that fitness in proliferation and survival in quiescence are poorly correlated in all three nutrient-restriction media (**Fig EV2A**). The fitness of the ∼3,900 mutants is distributed around 0 for each of the three proliferative conditions (**Fig 2B**), and the majority of mutants do not show a significant fitness defects compared to wild-type during proliferation (**Fig 2B & Fig EV2B**). By contrast, we find that many mutants show a survival defect in quiescent cells when starved for specific nutrients (**Fig EV2B**) resulting in increased variance in the distributions of survival compared to the distributions of fitness (**Fig 2B**). Critically, many of the genes that are dispensable for proliferative growth in each of the three media conditions are required for quiescence. For example, deletion of genes involved in the cAMP-PKA signaling pathway, GPB1/2, RGT2, GPR1 results in a profound survival defect in response to carbon starvation, but deletion of these genes does not impact fitness in carbon-restricted media (**Fig 2B left-panel**). Similarly, the autophagy genes *ATG4, ATG5, ATG7*, and *ATG12* have poor survival when starved for nitrogen, but do not have a fitness defect during proliferation in nitrogen-restricted media (**Fig 2B, mid-panel**). In response to phosphorus starvation, genes involved in response to pH have poor survival, but those same genes are dispensable for growth in phosphorus-restricted media (**Fig 2B right-panel**). Thus, the genetic requirements for growth in a specific nutrient-restricted media and induction of quiescence in response to starvation for that same nutrient are distinct.

**Figure 2.**
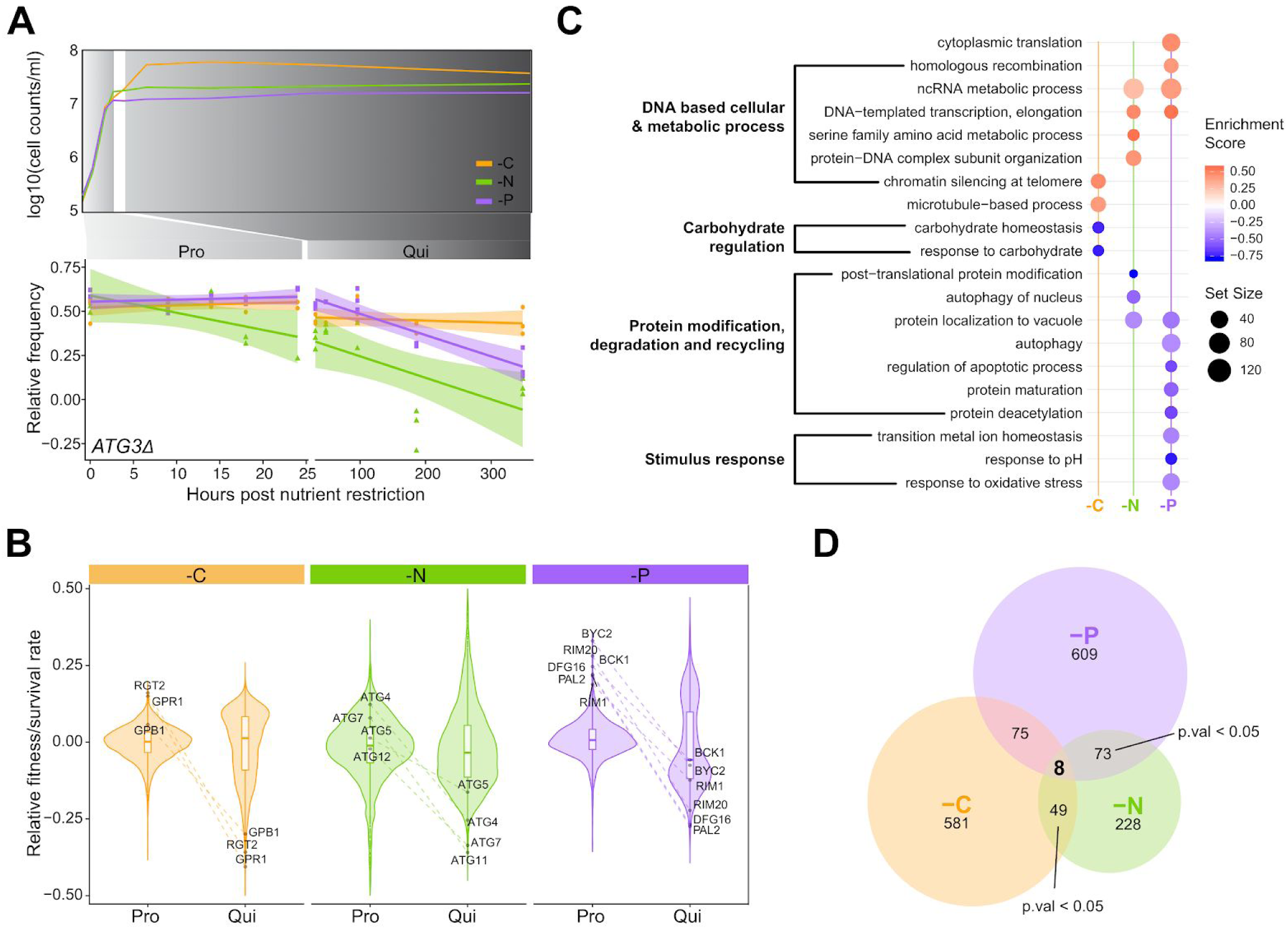
Distinct functional requirements for quiescence in response to different starvation signals. **A)** Cells exist in two distinct states depending on nutrient availability. An example of fitness, determined during proliferation, and survival, determined during quiescence, in the three different nutrient-restricted conditions is shown for *atg3Δ0*. **B)** Violin plot of the fitness and survival for each mutant during proliferation and quiescence in response to different nutrient restrictions. The indicated genes are examples of genes that are dispensable for proliferative growth in each of the three conditions but required for quiescence. **C)** Enriched GO terms identified using Gene Set Enrichment Analysis (GSEA). GSEA was applied to a ranked gene list based on the difference in survival during starvation and fitness during proliferation (S_Qui_ – F_Pro_) estimated using ANCOVA. The false discovery rate (FDR) was set at 0.05. Positive enrichment scores (red) indicate functions that have increased survival when starved (S_Qui_ – F_Pro_ > 0). Negative enrichment scores (blue) indicate functions that when impaired result in decreased survival (S_Qui_ – F_Pro_ < 0) during nutrient starvation. Set size indicates the gene number in each enriched term. **D)** Genes that are required for survival of starvation but dispensable for proliferation. We found 8 genes that are commonly required for survival of all three nutrient starvations (**Fig EV2D**); however, the overlap between conditions is not significant.

We identified gene functions that are specifically required for quiescence by performing Gene Set Enrichment Analysis (GSEA) (Yu et al. 2012; Subramanian et al. 2005) on gene lists ranked by the phenotypic difference between survival in quiescent conditions and fitness in proliferative conditions (S_qui_ – F_Pro_) (**methods**). We identified significantly enriched GO terms (Benjamini & Hochberg adjusted p-value < 0.05) and find that functions involved in responding to the specific starvation signal are required for survival. For example, mutants defective in carbon metabolism have reduced survival when starved for carbon, but the impairment of this function does not impact survival when starved for nitrogen or phosphorus (**Fig 2C**). Genes required for survival of nitrogen starvation are uniquely enriched for selective autophagy of nucleus related amino acid trafficking and recycling (**Fig 2C**). Some functional groups show similar requirements in response to both nitrogen and phosphorus starvation, such as autophagy and protein localization by the cytoplasm-to-vacuole targeting (CVT) pathway. By contrast, response to carbon starvation requires an entirely unique set of gene functions. Thus, the biological pathways and functions required for cellular quiescence differ between nutrient starvations.

### No evidence for common quiescence-specific genes

We sought to determine whether a common set of genes are required for quiescence in all starvation conditions. We identified a comparable number of quiescent-specific (hereafter: QS) genes detected in carbon (581) and phosphorus (609) restriction media. In nitrogen-restriction media, we identified about 2.5 times fewer QS genes: 228 (**Fig 2D**). To define a common set of QS genes, we applied three independent filtering criteria. We identified mutants that 1) are dispensable for proliferation in all three nutrient-restriction conditions (F_Pro_ >= 0, p.adj < 0.05), 2) show significant defects in quiescence in all three conditions (S_qui_ < 0,p.adj < 0.05), and 3) for which there is a significant negative difference between fitness and survival in all three conditions (S_qui_ – F_Pro_ < 0, p.adj < 0.05) (**Fig EV2C & methods**). Using these criteria, we found 8 genes that are commonly required for quiescence regardless of nutrient starvation (**Fig 2D & Fig EV2D**). However, this does not differ from what would be expected by chance. Thus, we find no evidence for the existence of a common set of QS genes that are required for establishing quiescence in response to carbon, nitrogen and phosphorus starvation.

### Detection of genetic interactions using pooled assays

We aimed to identify the set of genetic interactions between each non-essential gene and the three query kinase genes in three different nutritional conditions (carbon, nitrogen and phosphorus restricted media) and two different cellular states (proliferation and quiescence). As there have been limited studies using pooled fitness assays and time course data for quantifying genetic interactions, we considered two possible approaches for data analysis. First, we used analysis of covariance (ANCOVA) to compute the genetic interactions score (GIS) defined as the fitness (in proliferation) or survival (in quiescence) difference between the double (*queryΔ::kanMX xxx*^*n*^*Δ::natMX*) and single mutant (*hoΔ::kanMX xxx*^*n*^*Δ::natMX)* (**methods**). Briefly, in this case the two genotypes: single and double mutant are treated as independent variables in the model, scaled time is the covariate, and the normalized frequency at different timepoints are the dependent variable.

In a second approach, the genetic interaction score was calculated using the approach employed in previous genome-wide SGA which defines a null model based on a multiplicative hypothesis and defines a genetic interaction as a significant value of the observed and expected double mutant fitness: ε = *f* _*ab*_ − *f* _*a*_ · *f* _*b*_ (Costanzo et al. 2010). We computed the expected fitness for each double mutant by first computing the two single mutant fitness from the single deletion collection and then computing ε by determining the difference between the expected and measured fitness of double mutants. We find that the agreement between the two approaches is high (pearson’s *R* > 0.9) when applied to both fitness in proliferative cells and survival in quiescent cells (**Figure EV3A**). As ANCOVA has a well developed statistical framework for error estimation and significance testing, we elected to use ANCOVA to compute GIS for all subsequent analyses.

### Genetic interactions are condition dependent and common in quiescence

A genetic interaction is defined as a phenotypic effect in a double mutant that diverges from that expected on the basis of the same phenotype in each of the single mutants (Costanzo et al. 2010; Tong et al. 2004). We find that genetic interactions between genes are frequently condition dependent and differ both as a function of cellular state and environmental conditions. For example, in quiescent cells, the autophagy gene *ATG7* positively interacts with *TOR1* in carbon starvation, but negatively interacts with *TOR1* in phosphorus starvation (**Fig 3A** & **3B**). *ATG7* interacts negatively with *PHO85* and *RIM15* in phosphorus starvation but these interactions do not occur in carbon or nitrogen starvation (**Fig 3A** & **3B**). This example is illustrative of the conditional dependence of genetic interactions, which we find is the case for the vast majority of genotypes (the raw data and model fitting for all tested genetic interactions can be viewed in the associated web <u>application</u>).

**Figure 3.**
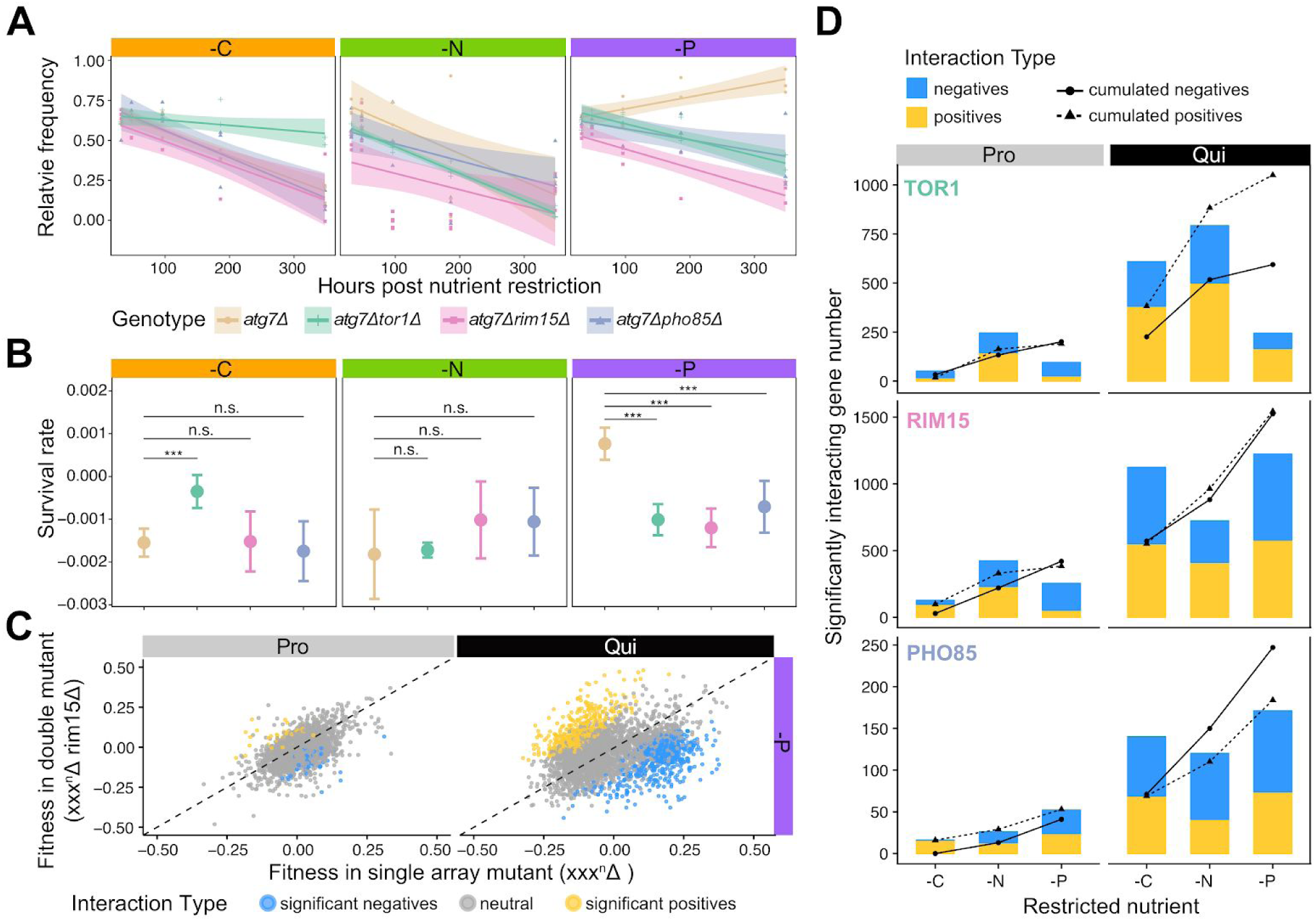
Identification of condition specific genetic interactions using pooled double mutant analysis. **A)** Genetic interactions for each gene were determined for three different query genes (*TOR1, RIM15*, and *PHO85*) in three different conditions (-C, -N, -P) and two different cellular states: quiescence (shown) and proliferation (not shown) using pooled mutant time series analysis. **B)** Survival rate for each genotype indicated in A) and the statistical significance estimated using ANCOVA. The significance is estimated when setting false discovery rate (FDR) at 5%. **C)** Relationship between single mutant phenotype (*xxx*^*n*^*Δ::natMX*) and the corresponding phenotype of the mutant in the background of a *RIM15* deletion (*rim15Δ::KanMX xxx*^*n*^*Δ::natMX*) in two different cellular states (Pro – proliferation, Qui – quiescence). The dashed line is the line of equality. Blue dots are genes that show a significant negative interaction with *RIM15* and yellow dots depict significant positive interactions. **D)** At a false discovery rate (FDR) of 5%, different numbers of significant genetic interactions are detected for three regulatory kinases in the three nutrient restrictions and two cellular states. Solid lines with circles indicate the cumulative total number of unique negative interactions and dashed lines with triangles indicate the cumulative total number of unique positive interactions.

Single mutants show stronger phenotypic defects in starvation conditions compared with growth conditions (**Fig 2B**). As single mutant fitness is predictive of genetic interaction propensity (Michael Costanzo et al. 2010), we find a weaker correlation between phenotypes of single (*hoΔ::kanMX xxxΔ::natMX)* and double (*rim15Δ::kanMX xxxΔ::natMX*) mutants in quiescent cells compared to proliferative cell (**Fig 3C, Fig EV3B**). More genetic interactions are detected in quiescent cells compared to proliferative cells regardless of the starvation signal (**Fig 3D**). For example, at an FDR of 5%, 55 genes show significant interactions with *TOR1* in proliferative cells growing in carbon-restricted media, whereas we identified 228 negative and 381 positive (∼6 times more) genetic interactions with *TOR1* in carbon-starved quiescent cells (**Fig EV3C**). This trend is observed for all three kinases (*TOR1, RIM15, PHO85*) in all starvation conditions (**Fig EV3C**). We detected both positive and negative interactions for each of the three kinases (**Fig 3D**) and an increase in total interactions for a given kinase as more conditions are assayed (**Fig 3D & Fig EV3D**) indicating that each additional assay reveals unique genetic interactions. We did not detect a bias in the number of positive or negative interactions in either cellular state.

### Genetic interaction profiles of kinases differ between cellular states

Genes that are functionally related tend to share a common set of genetic interactions that define a genetic interaction profile (Costanzo et al. 2010; Costanzo et al. 2016). As the activity of regulatory kinases depends on environmental signals, the functional consequences of deleting kinases is likely to be conditionally dependent, which may result in condition-dependent genetic interaction profiles. To identify the primary determinant of genetic interaction profiles in our study we quantified the similarity between all pairs of genetic interaction profiles for each kinase in each condition (C, N, P restricted conditions) and cellular state (proliferation and quiescence). Clustering of genetic interaction profiles reveals a clear distinction between proliferative and quiescent cells (**Fig EV4A**).

In quiescent cells, genetic interaction profiles of different kinases cluster as a function of the starvation signal (**Fig EV4A**) suggesting that the specific starvation signal is the primary determinant of the survival rate of double mutants regardless of the deleted kinase gene. By contrast, in proliferative conditions *TOR1, RIM15*, and *PHO85* genetic interactions profiles do not exclusively cluster by nutritional condition (**Fig EV4A**). These results indicate that genetic interaction profiles differ as a function of cellular state and that the impact of the environmental conditions on genetic interactions is variable.

To visualize the correlation between genetic interaction profiles for each kinase in each condition, we constructed correlation networks for both proliferative and quiescent cells (**Fig 4**). The correlation networks emphasize the importance of cellular state in determining the similarity of genetic interaction profiles as the genetic interaction profile similarity network is drastically remodeled in quiescence (**Fig 4B**) compared to proliferation (**Fig 4A**). For example, a negative correlation is detected between *TOR1* and *PHO85* in proliferative cells growing in carbon-restricted condition, but their genetic interaction profiles are positively correlated in carbon-starved quiescent cells (**Fig 4B** & **Fig EV4C**). For cells in the same physiological state, the environmental conditions can also alter the functional relationship between the same pair of kinases. For example, *RIM15* and *PHO85* genetic interaction profiles are highly correlated during growth in carbon-restricted media, but this similarity is greatly reduced during proliferation in nitrogen- and phosphorus-restricted conditions (**Fig 4A**). These results suggest that environmental conditions alter the regulatory relationships among signaling pathways both in quiescent and proliferative cells.

**Figure 4.**
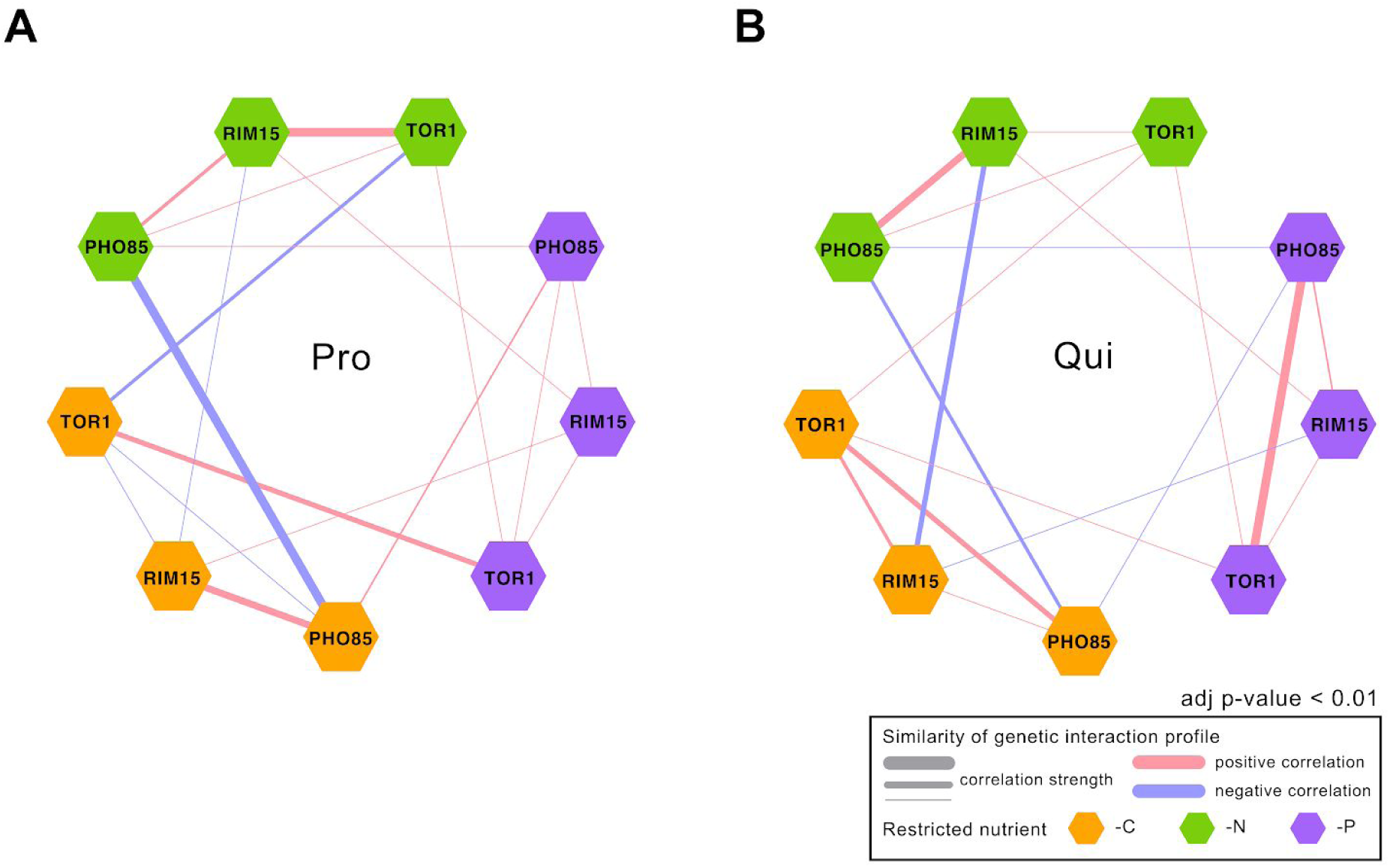
Genetic interaction profile similarities are condition dependent. **A)** Correlation networks based on genetic interaction profiles for *TOR1, RIM15*, and *PHO85* in proliferating cells (Pro) in three different nutrient restricted media: carbon (-C), nitrogen (-N), and phosphorus (-P). **B)** Correlation networks based on genetic interaction profiles for *TOR1, RIM15*, and *PHO85* in quiescent cells (Qui) induced by three nutrient starvations: -C, -N, -P. Hexagons are color coded based on the restricted nutrient type (orange for -C, green for -N and blue for -P). Kinases with positive pearson correlation are connected with pink edges and kinases with negative pearson correlation scores are connected with blue edges. The thickness of the edge indicates the strength of the correlation (i.e.. a larger absolute correlation is represented by thicker edge).

### Genetic interaction profiles are functionally coherent

To functionally annotate genetic interaction profiles for each kinase in each condition we used spatial analysis of functional enrichment (SAFE) (Baryshnikova 2016). SAFE maps quantitative attributes (i.e. the GIS) onto the reference network, defined by the correlation network of genome-wide genetic interaction profiles of 3,971 essential and non-essential genes, and tests for functional enrichment within densely connected regions, which define domains. Each of the 17 domains within this map comprises genes that share similar genetic interaction profiles and functional annotations (**Fig EV5**). We superimposed genetic interaction profiles of each kinase in each of the three nutrient-restricted media and both cellular states onto the reference network using SAFE. We find that kinases that show higher similarity in genetic interaction profiles (**Fig 4**) also show more similar enrichment patterns using SAFE analysis (**Fig 5**).

**Figure 5.**
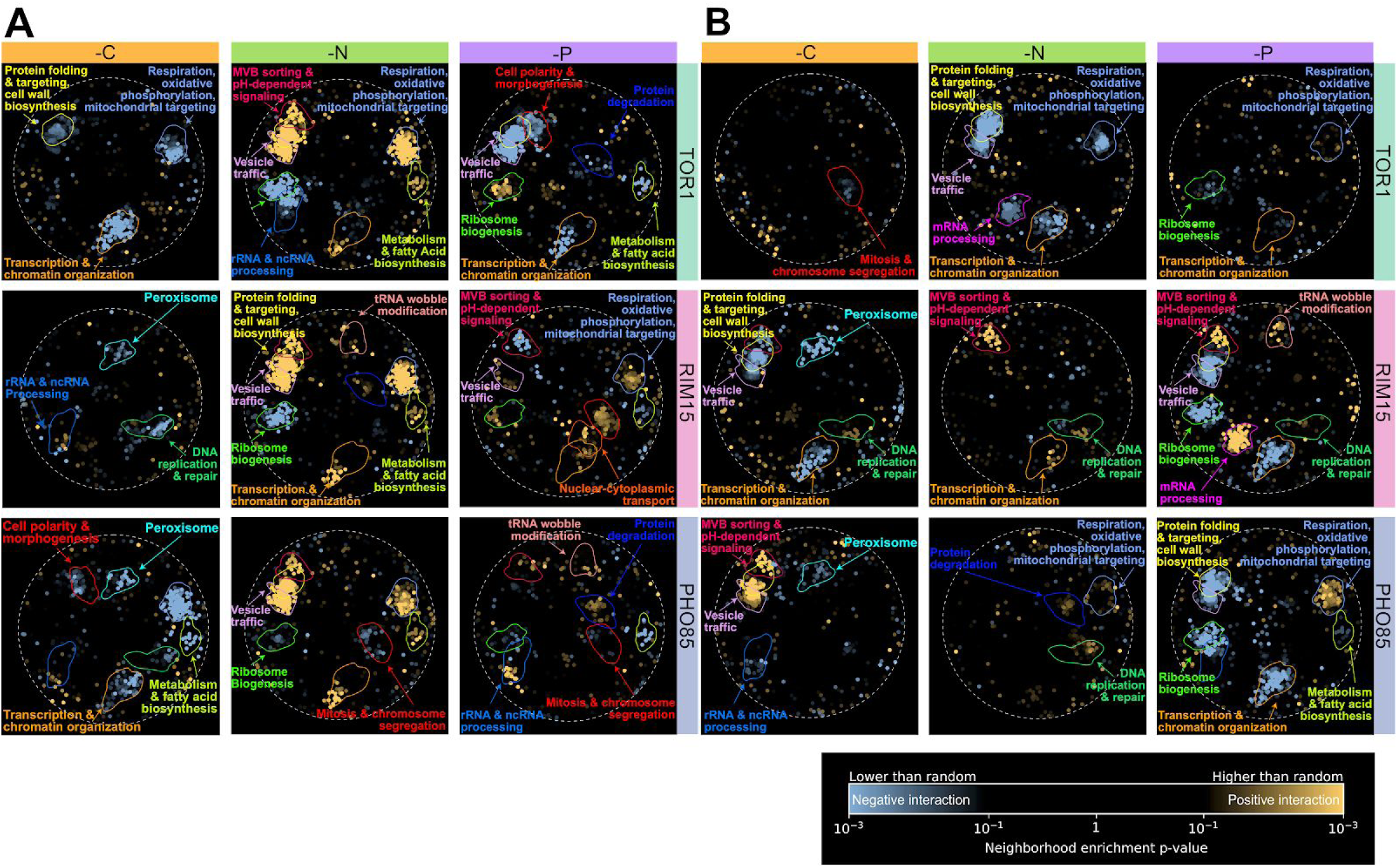
Functional mapping of kinase genetic interaction profiles in proliferating and quiescent cells. **A)** Genetic interaction enrichment landscape of *TOR1, RIM15, PHO85* in proliferating cells under different nutrient restrictions: carbon (-C), nitrogen (-N), phosphorus (-P). **B)** Genetic interaction enrichment landscape of *TOR1, RIM15, PHO85* in quiescent cells in response to different nutrient starvations (-C, -N, -P). Each dot represents one gene. Blue dots represent genes have negative interactions with corresponding kinase (row-wise) in each condition (column-wise), and yellow dots represent genes with positive interactions.

The functional annotation of genetic interactions for each kinase differs as a function of the cellular state. For example, functional domains related to respiration, oxidative phosphorylation, mitochondrial targeting, transcription, and chromatin organization are enriched for negative genetic interactions with *TOR1* and *PHO85* in carbon restricted proliferative cells (**Fig 5A**), but we find no evidence for enrichment in quiescent cells starved for carbon (**Fig 5B**). Similarly, in nitrogen restricted conditions, *TOR1, RIM15* and *PHO85* share similar coherent functional interactions in proliferative cells, which are not observed in quiescent cells starved for nitrogen.

In addition, the functional enrichment of genetic interactions for each kinase differs between the three different nutrient-restricted conditions. For example, ribosome biogenesis genes are enriched for negative interactions with *TOR1* in nitrogen-restricted proliferative cells (**Fig 5A**), but in phosphorus-restricted proliferative cells ribosome biogenesis genes positively interact with *TOR1*. We find multiple additional cases of enrichment within functional domains, in which the sign of the genetic interactions is opposite between nitrogen and phosphorus restrictions in *TOR1* (**Fig 5A**), suggesting that *TOR1* may play different regulatory roles in responding to nitrogen and phosphorus restriction.

We have also found cases of functional enrichment that are maintained in the two different cellular states. For example, genes involved in peroxisome functions are enriched for negative interactions with *RIM15* and *PHO85* in carbon restricted proliferative cells and carbon starved quiescent cells (**Fig 5A**, cyan arrow/circle). This suggests that in carbon-restricted conditions, *RIM15* and *PHO85* may function within the same or similar pathway to maintain long chain fatty-acid recycling and provide energy for cells in calorie-restricted conditions.

### Common and specific genetic interactions with *RIM15* support its role as a central mediator of quiescence

RIM15 has previously been identified as an integrator of quiescence signals that is downstream of TOR1, PHO85 and PKA (Pedruzzi et al. 2003; Wanke et al. 2005; Olivares-Marin et al. 2018). Therefore, we might expect that the genetic interaction profiles for *RIM15* should show more functional coherence in response to different quiescence signals compared to *TOR1* and *PHO85*, which are upstream of RIM15. As the reference genetic interaction map used for SAFE does not include all genes in our genetic interaction dataset (only ∼2,900 non-essential genes are present in the reference), we applied gene set enrichment analysis to the list of genes that significantly interact with each kinase (**method and materials**). Due to the limited number of significant interactions detected in proliferative cells (**Fig 3D** and **Fig EV3C**), we do not find any enriched GO terms for any kinase. However, we identified multiple significantly enriched functional categories in quiescent cells. As with SAFE analysis, the functional enrichment of the significant interacting genes for a given kinase depends on the starvation signal (**Fig 6A**).

**Figure 6.**
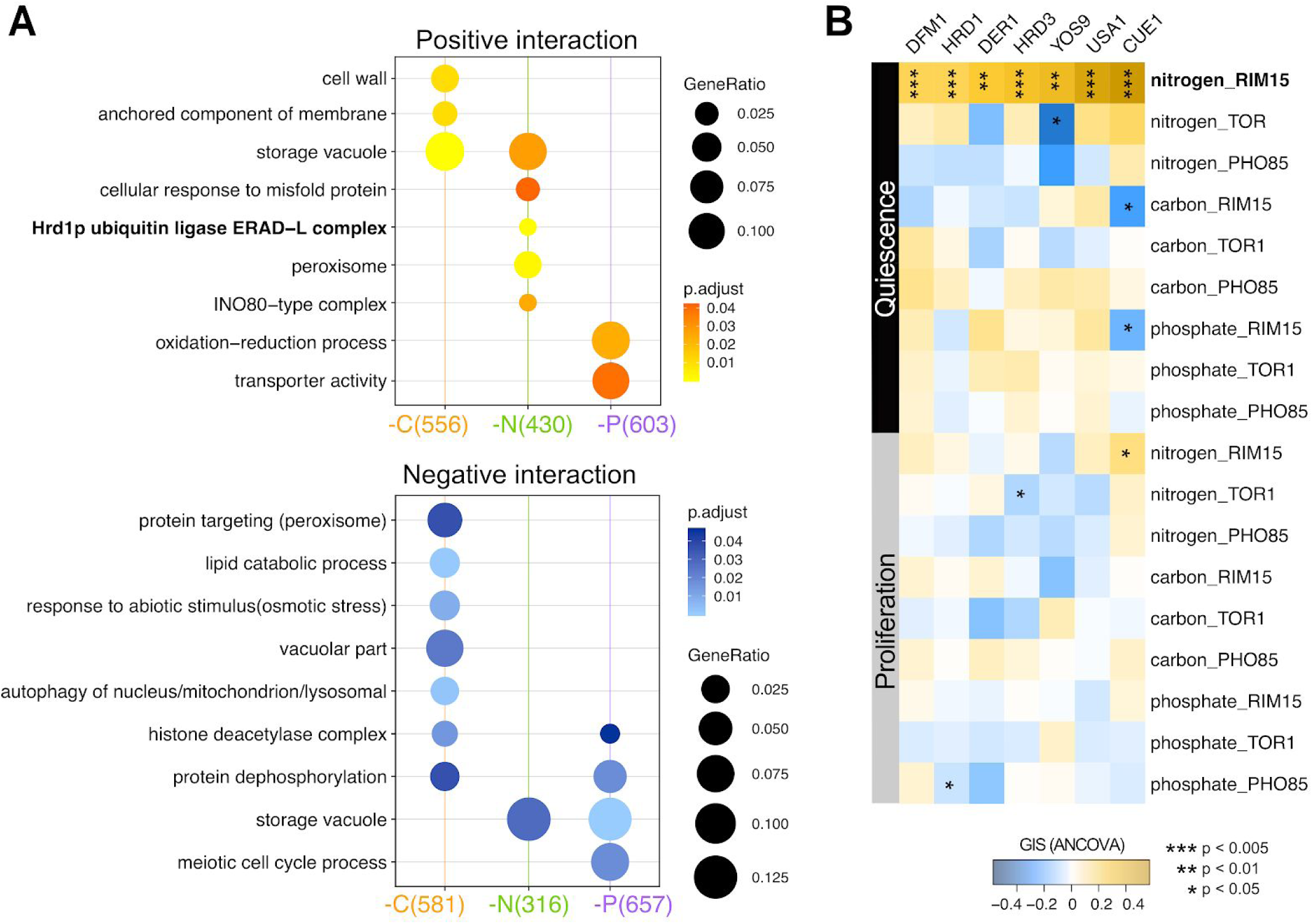
*RIM15* genetic interactions profiles indicates it is an integrator of quiescence signals with nutrient-specific functions. **A)** GO term enrichment analysis for genes that significantly interact with *RIM15* in all nutrient starvation conditions. Only significantly enriched GO terms are shown (p.adjust < 0.05). (yellow – positive interaction, blue – negative interaction). **B)** Genetic interaction profile of the genes encoding the ERAD-L complex. ERAD-L genes show a unique cohesive set of positive genetic interactions with *RIM15* in nitrogen starvation-induced quiescent cells. Each column is the genetic interaction score between ERAD-L genes and *RIM15* quantified using ANCOVA, and each row is the genetic interaction score between ERAD-L genes and each of the other kinases in each nutrient restricted condition.

*RIM15* shows more common genetic interactions in response to different starvation signals in comparison with the other two kinases. Three functional groups are shared among genes interacting with *RIM15* in response to carbon/nitrogen or nitrogen/phosphorus starvations (**Fig 6A**, lower panel) whereas there is no functional overlap detected for *TOR1* or *PHO85* genetic interaction profiles under the same conditions (**Fig EV6A**). This is consistent with a model in which RIM15 regulates quiescence through integration of diverse signals and execution of similar regulatory interactions. In quiescent cells, *RIM15* shown consistent negative genetic interactions with genes involved in vacuolar functions regardless of the starvation signals perhaps reflecting a role for *RIM15* in regulating autophagy and protein recycling in response to different starvations.

Interestingly, we find that genes that function in the Endoplasmic-reticulum-associated protein degradation, luminal domain monitored (ERAD-L) pathway show coherent positive interactions with *RIM15* specifically in nitrogen-starved quiescent cells (**Fig 6A**). This Includes each of the genes that is known to function in ERAD-L: *USA1, YOS9, DFM1, HRD1, HRD3, CUE1, and DER1* (**Fig 6B** and **Fig EV6B**). ERAD-L genes present in the genetic interaction reference data used for SAFE analysis; *HRD1, HRD3, CUE1, and USA1* are found in the domain enriched for ubiquitin-dependent protein catabolic process (**Figure EV6C**, red arrow) pointing to a specific function for RIM15 in proteostasis regulation in response to nitrogen starvation.

## Discussion

Cellular quiescence is the predominant state of eukaryotic cells. To study the genetic requirements of cellular quiescence in yeast cells we quantified the effect of each gene deletion in response to three distinct nutrient starvation signals (carbon, nitrogen, phosphorus). To study how these signals are coordinated within quiescent cells we quantified genetic interactions with three regulatory kinases in each of the three starvation conditions. To undertake this study we quantified phenotypic differences in different cellular states (proliferation versus quiescence) and genotypes (single versus double mutational background) using multiplexed barcoded analysis to track thousands of different genotypes using time course analysis. By testing the contribution of ∼3,900 yeast non-essential genes to fitness in proliferating cells and survival in quiescent cells in three different nutrient-restricted conditions we find no evidence for genes that are commonly required for quiescence. We extended our method for multiplexed analysis of genotypes to study ∼11,700 double mutants encompassing three core kinases: TOR1, RIM15 and PHO85, which allowed us for the first time to test genome-wide for genetic interactions with regulatory kinases that mediate quiescence.

### Distinct gene functions are required for quiescence in response to different nutrient starvations

The functional requirements for maintaining and exiting quiescence differ depending on starvation signals. Time course analysis of fitness during proliferation and survival during starvation (**Fig 2C**) support previous findings that yeast cells have distinct functional requirements for maintaining viability of quiescent cells in response to different nutritional starvations (Gresham et al. 2010; Klosinska et al. 2011). In addition, our results show that a substantial fraction of the non-essential yeast genome is required for survival during quiescence independent of their requirements for growth. For example, in carbon-restricted conditions, deletion of 713 (∼15%) of the non-essential genes results in a significant defect in quiescence (**Fig 2D**). Clearly, the definition of an “essential gene” is dependent on the condition in which essentiality is assessed.

Across all starvation conditions, we found that only 8 genes are commonly required for quiescence, a result that is not significantly different from chance (**Fig 2D**). The absence of a common set of genetic requirements for quiescence in response to different nutrient starvations is consistent with earlier work (Klosinska et al. 2011). Although there appears to be no common set of requirements for quiescence, we do find that different nutrient starvations share genetic requirements for quiescence. Nitrogen- and phosphorus-starved quiescent cells tend to have more overlapping features compared to carbon-starvation induced quiescence. For example, 81 genes are commonly required for maintaining quiescence in response to nitrogen or phosphorus starvation, whereas only 57 genes are commonly required for quiescence in nitrogen and carbon starvation (**Fig 2D**). Results from functional enrichment analysis are consistent with the trend of greater overlap in genetic requirements in nitrogen and phosphorus starvation. For example, genes involved in protein localization by CVT pathway are required in response to nitrogen or phosphorus starvation. The lack of commonly required functions for response to nitrogen, phosphorus, and carbon starvations may reflect the different primary biological uses of carbon, nitrogen and phosphorus: carbon is the major energy source, whereas nitrogen and phosphorus are primarily required for macromolecular synthesis (de Virgilio 2012; Broach 2012; Alberts et al. 2013; Wilson and Roach 2002).

### Expanding phenotypic space to identify novel genetic interactions

To date, genome-wide genetic interaction mapping has primarily been performed in a single condition and assayed using a single phenotype – growth in rich media. Our genome-wide genetic interaction mapping in different conditions and cellular states indicates that: 1) genetic interactions with regulatory kinases vary between conditions; 2) genome-wide genetic interaction mapping is extensible to additional phenotypes; and that 3) for a given physiological state, increasing the number of conditions results in an increase in the number of significant GIs. This latter point is consistent with a recent study that investigated genetic interactions in different growth conditions (Jaffe et al. 2019). Despite the fact that our genetic interaction data set is limited in its scale and is focused on regulatory kinase genes, we anticipate that our methodology can be broadly applied to define genetic interactions in different conditions and cellular states.

### Novel function of RIM15 in autophagy and ERAD-L

Endoplasmic-reticulum-associated protein degradation (ERAD) is a quality control mechanism that ensures only properly folded proteins leave the ER. Autophagy has been proposed to be a backup mechanism for ERAD. Previous study has shown that RIM15 plays a role in regulating autophagy and protein homeostasis (Waliullah et al. 2017; Huang et al. 2018). In our study we find that genes that function in ERAD show coherent positive interactions with RIM15 in nitrogen starvation conditions, suggesting that RIM15 regulation of ERAD activity in response to nitrogen starvation is essential for quiescence. It is possible that RIM15 functions to regulate clearance of stress-induced misfolded proteins during nitrogen starvation by mediating the balance between autophagy and ERAD.

### Implications for quantitative genetics

Our study has important implications for our understanding of the genotype to phenotype map. The prevailing result from our study is that the effect of a given gene deletion on a phenotype (either fitness or survival) is highly dependent on the specific environmental conditions of the cell. Although nitrogen, carbon and phosphorus starvation all lead to cell cycle arrest and the initiation of quiescence, the genetic requirements for this behavior are distinct. We find that the conditional dependence extends to genetic interactions as we detect different sets of genetic interactions in different growth and starvation conditions. These results are consistent with our previous study of natural genetic variation in which we found that the effect sizes of QTL underlying fitness differences, and genetic interactions between QTL, are acutely sensitive to the composition of the growth media (Ziv et al. 2017). Identifying quantitative genetic effects and interactions that are insensitive to environmental variation appears challenging and may, in fact, be extremely rare.

### Implications for the study of cellular quiescence in yeast

It has been argued that starvation for glucose is the relevant condition for studying quiescence (Sagot and Laporte 2019) and indeed the vast majority of quiescence studies are performed in conditions in which carbon starvation is the pro-quiescence signal (Laporte et al. 2011; Laporte et al. 2018). However, it has been appreciated for many decades that yeast cells can initiate a quiescence state in response to different starvation signals (Lillie and Pringle 1980). Our study reiterates the importance of studying quiescence in response to different nutrient starvation conditions. Many important biological processes are likely to be missed – autophagy being a preeminent example – if carbon starvation is the only condition studied. Organisms in the natural world experience a range of nutrient limitations and nitrogen and phosphorus appears to be the predominant limiting nutrients in most ecologies (Elser et al. 2007). Thus, a complete understanding of cellular quiescence requires the study of different nutrient starvation signals.

### Relevance to aging and cancer

The study of cellular quiescence may inform our understanding of cellular aging and provide insight into the therapeutic challenge of dormant cancer cells. Our study supports previous findings that quiescence establishment does not follow the same route depending on the nature of the inducing signal (Coller et al. 2006; Klosinska et al. 2011). In addition, different ‘degrees’ of quiescence may exist (Gookin et al. 2017; Coller et al. 2006; Laporte et al. 2017) as we find that cells maintained longer in quiescence need more time to return to growth (data not shown). Thus, quiescence may be viewed as a continuum that ultimately leads to senescence (even if that may take thousands of years) unless conditions favorable for proliferation are met.

Overall, our data highlight the fact that quiescence does not imply uniformity (O’Farrell 2011). The idea that quiescence establishment is the result of a universal program is clearly an over-simplification. Our study points to a rich spectrum of condition-specific genetic interactions that underlie cellular fitness and survival across a diversity of conditions and introduces a generalizable framework for extending genome-wide genetic interaction mapping to diverse conditions and phenotypes. Deciphering the underlying regulatory rationale and the hierarchical relationships between these signaling pathways in different conditions is critical for understanding cellular quiescence.

## Materials and Methods

### SGA Library construction

The haploid prototrophic double deletion collections were constructed using the synthetic genetic array method (Tong et al, 2001). The genotype and ploidy of double mutants are prototrophic haploid (**Fig EV1A**). For the single deletion collection (array mutants), gene deletion alleles are marked with the kanMX4 cassette conferring G418 resistance, which is flanked by two unique molecular barcodes (the UPTAG and DNTAG). For double deletion collection, an additional query allele is marked with NatR cassette conferring nourseothricin resistance. To construct the RIM15 and TOR1 SGA query strains we mated a MATa xxxΔ0::NATr strain (transformed from FY4 with a NATr PCR product targeting the xxx allele) with the Y7092 strain. A haploid prototrophic strain was identified following tetrad dissection and genotyping using selective media with G418 and nourseothricin. To construct the HO, and PHO85 SGA query strains we transformed a prototrophic strain containing the SGA marker with a NATr PCR product targeting the xxx allele. Insertion of NATr was confirmed via PCR and the genotype of the strain was checked via replica plating onto selective media resulting in strains listed in **Table EV1**.

### Growth conditions

After growth of individual selected mutants on YPD agar plates, all mutants were pooled to a final density around 1.7 × 10^9^ cells/ml. Each agar plate contained single colonies of individual genotypes and replicated colonies of the control *hoΔ* strain. We inoculated 1.5 × 10^8^ cells into 300ml of nutrient limited medium: for glucose- (C, 4.4mM carbon), ammonia- (N, 0.8mM nitrogen), and phosphorus- (P, 0.04mM phosphorus) at 300ml. To define the fitness of ∼ 4,700 mutants within each nutrient limiting conditions and growing stage, we performed three independent experiments for each mutant per nutrient limiting conditions. In total, we had 4 mutant collections × 3 biological replicates × 3 nutrient limiting conditions in bioreactors used to maintain the temperature at 30 degrees and pH at 5. To model the fitness of each genotype at different states spanning both proliferative and quiescence stages, we collected five time points in each stage (based on growth curve (**Figure EV1B**). The duration of the experiment was 15∼16 days, and populations were sampled at 0, 9, 14, 18, 24, 32, 48, 96, 187, 368 hours for outgrowth and barcode sequencing. To isolate viable cells from the stationary phase culture, we transferred 1mL (i.e., 2 × 10^5^ cells) from the pooled library at each time point into 5 mL minimal cultures. Cells were grown for 24∼32 hr to a final density of 3 × 10^8^ cells/mL in all conditions. Cells were then washed with water once, and then resuspended in 1mL sorbitol buffer for genomic DNA purification.

### DNA extraction and library preparation for Bar-seq

Genomic DNA was isolated from 1 × 10^8^ cells for each sample (3 nutrient-restriction × 3 biological replicates × 4 deletion collections × 10 times points) using invitrogen PureLink™ Pro 96 Genomic DNA Purification Kit. We adapted the two-step PCR protocol for efficient multiplexing of Bar-seq libraries (David G. Robinson *et al*, 2013). Briefly, UPTAGs and DNTAGs were amplified separately from the same genomic DNA template. In the first PCR step, a unique sample indices are added to each sample. For each biological replicate, we used 120 unique sample indices that differed by at least two nucleotides to label each sample from 3 nutrient limiting conditions × 4 deletion collections × 10 timepoints. We normalized genomic DNA concentrations to 10 ng/ml and used 100 ng template amplified barcodes using the following PCR program: 2 min at 98°C followed by 20 cycles of 10 sec at 98°C, 10 sec at 50°C,10 sec at 72°C, and a final extension step of 2 min at 72°C. PCR products were confirmed on 2% agarose gels and quantified the concentration using a SYBRGreen staining followed by Tecan Freedom Evo and Infinite Microplate Reader. We combined 35 ng from each of the 120 different UPTAG libraries and, in a separate tube, 35 ng from each of the 120 different DNTAG libraries for each condition/deletion collection. The multiplexed UPTAG libraries were then amplified using the primers P5 (59-A ATG ATA CGG CGA CCA CCG AGA TCT ACA CTC TTT CCC TAC ACG ACG CTC TTC CGA TCT-39) and Illumina_UPkanMX, and the combined DNTAG libraries were amplified using the P5 and Illumina_DNkanMX primers using the identical PCR program as the first step with 100 ng template. The 140-bp UPTAG and DNTAG libraries were purified using QIAquick PCR purification columns, quantified using a Qubit fluorometer for qPCR quantification, combined in equimolar amounts after qPCR, and adjusted to a final concentration of 4 nM mixture of pooled UPTAG and DNTAG. In total, each sequencing library contained 120 UPTAG and 120 DNTAG libraries from 120 different samples. The library was sequenced on a single lane of an Illumina NextSeq 500 with HighOutput 1 × 75bp read configuration. 20% PhiX was spiked into each library for increasing the complexity of two color base calling on Illumina NextSeq500 platform.

### Data analysis, filtering and normalization

Sequence reads were matched to the yeast deletion collection barcodes using re-annotated by Smith *et al*. (2009). Inexact matching was performed by identifying barcode sequences that were within a Levenshtein distance of 2 from each read (Levenshtein 1966). Sample indices were similarly matched using a maximum Levenshtein distance of 1. The final matrix of counts matching the UPTAG and DNTAG of each of the 360 samples is provided as **Table EV3**. 52 libraries with total read depth less than 1 × 10^5^ reads were removed from the 720 libraries. We merged the UPtag and DOWNtag counts representing the same gene within each condition resulting in 311 libraries in total. A set of outliers was identified that had fewer than 3,000 total reads across all 311 samples. These low-count matches were likely due to sequencing error and were removed. 1,996 mutants were removed with a coverage less than 3,000 or missing in either tag counts. After filtering, a matrix containing 3,931 mutants consistent with high quality counts data across 311 conditions was generated corresponding to 692,755,604 sequence reads. This counts table was normalized using the function varianceStabilizingTransformation in the DESeq2 package (Love et al, 2014) (version 1.8.1) with arguments blind = FALSE and fitType = “local”.

### Fitness, survival, and phenotypic difference quantification

The normalized frequency of each mutant within each library were used for linear regression modeling. For example, in HO library, the count for each mutant (*ho::kanMX xxx*_*n*_::*natMX*) is normalized by the count for the wild type control (*hoΔ::kanMX his3Δ1 can1Δ::STE2pr-Sp_his5)* at corresponding time points. In the other double mutant libraries, the counts for each double mutant (*query::kanMX xxx*_*n*_::*natMX*) is normalized by the counts of the query mutant (*queryΔ::kanMX his3Δ1 can1Δ::STE2pr-Sp_his5)* at corresponding time points. For each mutant strain *N*, fitness *f* _*n*_ was calculated as the coefficients of linear regression model calculated in R:

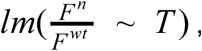

therefore,

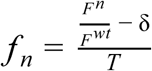

with *F*^*n*^ being the normalized counts of strain *N* at each time point and *F*^*wt*^ is the normalized counts of pseudo wild type strain at each time point. *T* refers to timepoints, which was measured in days for quantifying the fitness in prolonged starvation. δ is the error term.

In order to compare the phenotypic difference for a given mutants between different cellular states, before building linear regression model for each mutant in proliferation or quiescence, we scaled the independent variable, time (hours) for each stage into the same unit but maintaining the natural interval using *scale*() function in R. For example, the time point (independent variable) in proliferative stage were scaled from 0h, 9h, 14h, 18h, 24h into 0, 0.5246676, 0.8161497, 1.0493353, 1.3991137, and the time point for sample collected during quiescence were scaled from 32h, 48h, 96h, 187h, 368h into 0.1553874, 0.2330811, 0.4661622, 0.9031894, 1.6995499. The, we are able to quantify the phenotypic difference between fitness in proliferation and survival in quiescence using ANCOVA for a given mutant:

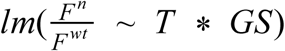

Where *T* is the scaled time and *GS* is the <u>G</u>rowing <u>S</u>tage (e.g. proliferation vs quiescence). The different growth stages in this function is the interaction term, which was tested for statistical significance.

After quantifying the fitness difference between quiescence and proliferation for a given mutant, we ranked the mutants by fitness difference in a descending order and then applied gene set enrichment analysis (GSEA) of ranked genes using clusterprofiler (Yu et al. 2012).

To understand whether the common genes that required in response to different quiescent signals are statistically significant or not, we implemented the proposed multi-set intersection test algorithm in an R software package *SuperExactTest* (Wang et al. 2015). *This framework is used to* compute the statistical distributions of multi-set intersections based upon combinatorial theory and accordingly designed a procedure to efficiently calculate the exact probability of multi-set intersections. The inputs for *SuperExactTest* include three lists of genes that are essential in quiescence. The three lists corresponding to three conditions (-carbon, -nitrogen, -phosphorus) and the size of the background population from which the sets are sampled is 5,927.

### Comparison of SGA genetic interaction quantification with ANCOVA

#### SGA genetic interactions scoring method

We first computed genetic interactions using a method analogous to estimation of epsilon (ε) as defined in classical SGA screens from the Boone lab. The SGA-like score was quantified by testing the null hypothesis based on a multiplicative model from single mutant fitness :

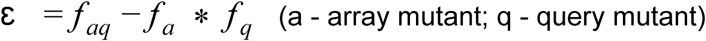

In our case, ε is calculated as the difference between the coefficients of linear modeling:

where

Therefore,

*f*_*aq*_ is the coefficients generated by 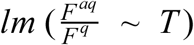,

*f*_*a*_ is the coefficients generated by 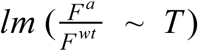,

*f*_*q*_ is the coefficients generated by 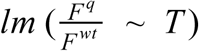,

Therefore *f*_*aq*_, *f*_*a*_, *f*_*q*_ should be normally distributed around 0 with positive (better than WT) and negative (worse than WT) fitness. To estimate the expected fitness in double mutant based on multiplicative model, we take the *exp*() of the coefficients for each model to eliminate the discordance of the signs in fitness. Then we calculated the expected fitness using multiplicative model:

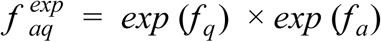

Therefore

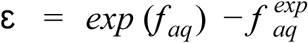

The standard error *S*_*a*_ & *S*_*q*_ are the standard error of each linear model. The standard error in expected fitness is calculated by propagating standard error from each individual model:

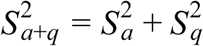

Then, the statistical significance between expected (multiplicative model) and observed model was calculated by *Welch’s t-test*:

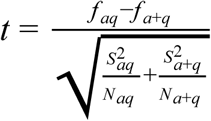

where the degrees of freedom associated with this variance estimate is approximated using the Welch-Satterthwaite equation:

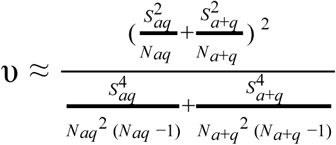

#### Genetic interactions quantification by ANCOVA

All libraries were normalized by the common query deletion. Therefore, our GIS can be calculated by looking at the difference between normalized fitness without worrying about the query mutant phenotype,

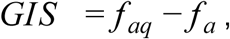

Where

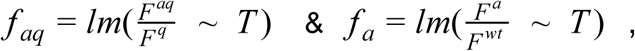

In this case, the genetic interaction is calculated directly by testing whether the query mutation significantly changes the relationship between time and relative fitness for a given mutant. We applied ANCOVA using:

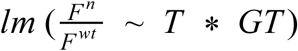

Where *T* is the scaled time and *GT* is the <u>G</u>eno<u>T</u>ype (e.g.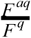 vs 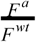). The significance of the interaction term was determined using a standard *t test*.

### Functional annotation with clusterProfiler

Gene Set Enrichment Analysis (GSEA) was applied on the ranked gene list based on phenotypic difference using clusterProfiler (Yu et al. 2012). GO overrepresentation test was applied to significantly interacting genes and quiescent specific gene list (Table EV13-18).

### Network Construction using Cytoscape 3.0

The correlation among genetic interaction profiles were calculated by metScape 3 Correlation Calculator v1.0.1 using DSPC method and then visualized in Cytoscape 3.0.

### Spatial Analysis of Functional Enrichment (SAFE)

The systematic functional annotation and visualization of interaction profile for all kinase under different conditions and cellular states was applied without any filtering on the interaction list. In this enrichment analysis we used all genes without filtering based on statistical interaction significance (from ANCOVA). This is because isolated false positives are scattered throughout the entire network, which do not typically result in significant enrichment anywhere in the network. Meanwhile, weak but consistent effects, e.g. genes having weaker or less significant GIs but clustering together in the network are very interesting.Thevisualizationandlocalenrichmentannotationwasperformed using SAFE (Baryshnikova 2016).

## Supporting information

Sun et al Table S3

Sun et al Table S4

Sun et al Table S5

Sun et al Table S6

Sun et al Table S7

Sun et al Table S8

Sun et al Table S9

Sun et al Table S11

Sun et al Table S12

Sun et al Table S13

Sun et al Table S14

Sun et al Table S15

Sun et al Table S16

Sun et al Table S17

Sun et al Table S18

## Acknowledgements

We thank Charlie Boone and Michael Costanzo for library construction. We thank the members of the Gresham lab and Vogel lab for helpful discussions. We thank NYU Gencore for next generation sequencing and FACS, This work was supported by grants from the NIH (R01 GM107466) and NSF (MCB1818234).

## Authorship contributions

DG and SS designed the study; SS and NB performed the experiments; SS analyzed the data; and DG and SS wrote the manuscript.

## Data availability and analysis scripts

Sequencing counts and normalized relative frequencies, as well as fitness, survival, interaction score estimates for each nutrient-restricted or starved condition can be found on OSF (https://osf.io/6avpn/). The custom Python and R scripts used to parse raw sequencing data, analyze results, build the shiny web-app and generate manuscript figures are available on GitHub (https://github.com/ss6025/GI-of-kinases-in-quiescence_2018). All quantifiable genes can be queried interactively using the shiny app available at: http://shiny.bio.nyu.edu/ss6025/shiny_Genetic_Interaction/. The raw FASTQ files from Illumina sequencing are publically available under the NCBI BioProject under Accession Number PRJNA559194.

## Expanded View tables

**Table EV1.** List of strains used and generated in this study.

| Strain ID | Gene/Alias | Genotype | Notes |
| --- | --- | --- | --- |
| Background Strain | NA | MATα can1Δ::STE2pr-Sp_his5 lyp1Δ his3Δ1 | Prototrophic strain background |
| YJR066W_y13791 | TOR1 | - MAT $\alpha$ tor1ΔNatR can1Δ::STE2pr-Sp_his5 lyp1Δ his3Δ1 LYS2+ URA3+ MET3+ LEU2+ | Prototrophic SGA query strain |
| YFL033C_y13792 | RIM15 | - MAT $\alpha$ rim15ΔNatR can1Δ::STE2pr-Sp_his5 lyp1Δ his3Δ1 LYS2+ URA3+ MET3+ LEU2 + | Prototrophic SGA query strain |
| YDL227C_y13819 | HO | - MAT $\alpha$ hoΔ::NatR can1Δ::STE2pr-Sp_his5 lyp1Δ his3Δ1 LEU2+ URA3+ MET15+ LYS2+ | Prototrophic SGA control strain |
| YPL031C_y13820 | PHO85 | - MAT $\alpha$ pho85Δ::NatR can1Δ::STE2pr-Sp_his5 lyp1Δhis3-1 LEU2+ URA3+ MET15+ LYS2+ | Prototrophic SGA query strain |

**Table EV2.** Media used in this study.

Carbon restricted media (per 1L)
| Compound | Carbon restricted | Carbon supplemented |
| --- | --- | --- |
| 40% Glucose | 2.0 mL | 50 mL |
| 1000X vitamin | 1 mL | 1 mL |
| 1000X metals | 1 mL | 1 mL |
| 10X carbon salts | 10 mL | 10 mL |
| **Final (C) concentration** | **0.08%** | **2%** |

Nitrogen restricted media (per 1L)
| Compound | Nitrogen restricted | Nitrogen supplemented |
| --- | --- | --- |
| Glucose | 20 g | 20 g |
| 1000X vitamin | 1 mL | 1 mL |
| 1000X metals | 1 mL | 1 mL |
| 50 mM (NH<sub>4</sub>)<sub>2</sub>SO<sub>4</sub> (solution) | 8 mL |  |
| 10X nitrogen salts | 10 mL | 10 mL |
| **Final (N) concentration** | 800μM | 38mM |

Phosphorus restricted media (per 1L)
| Compound | Phosphorus restricted | Phosphorus supplemented |
| --- | --- | --- |
| Glucose | 20 g | 20 g |
| 1000X vitamin | 1 mL | 1 mL |
| 1000X metals | 1 mL | 1 mL |
| 10g/L KH<sub>2</sub>PO<sub>4</sub> (solution) | 0.5 mL | 100 mL |
| 10X phosphorus salts | 10 mL | 10 mL |
| **Final (P) concentration** | 5 mg/L | 1 g/L |

## Expanded View table legends

Table EV3: Filtered and variable stabilization transformed counts table. Original data for Fig 1C & Fig 2A.

Table EV4: Linear regression modeling of mutant fitness over prolonged starvation for each replicate. Original data for Fig 1B.

Table EV5: Linear regression modeling of mutant fitness over prolonged starvation across all replicates. Original data for Fig 1C.

Table EV6: Fitness and survival rate modeling for cells in different growth stages. Original data for Fig2B & FigS2A.

Table EV7: Phenotypic difference between quiescence and proliferation quantified by ANCOVA.

Table EV8: Fitness, survival rate and the difference in phenotype values.

Table EV9: Quiescent Specific (QS) genes in different starvation conditions. Original data for Fig 2C.

Table EV10: Core set of genes required for quiescence across different starvation conditions. Original data for Fig S2D

Table EV11: Gene Set Enrichment Analysis results for QS genes in three different conditions.

Table EV12: Genetic interaction scores quantified by ANCOVA using different phenotypic readouts for proliferation and quiescence.

Table EV13: Genetic interaction profiles for all kinases in all conditions during proliferation. Original data for Fig 4A.

Table EV14: Genetic interaction profiles for all kinases in all conditions during quiescence. Original data for Fig 4B.

Table EV15: Functional annotation of gene clusters that positively interact with RIM15 in quiescent cells. Original data for Fig 6A.

Table EV16: Functional annotation of gene clusters that negatively interact with RIM15 in quiescent cells. Original data for Fig 6B.

Table EV17: Functional annotation of gene clusters that positively interact with TOR1 in quiescent cells. Original data for Fig 6SA.

Table EV18: Functional annotation of gene clusters that negatively interact with TOR1 in quiescent cells. Original data for Fig 6SB.

## Expanded View figures

**Figure EV1.**
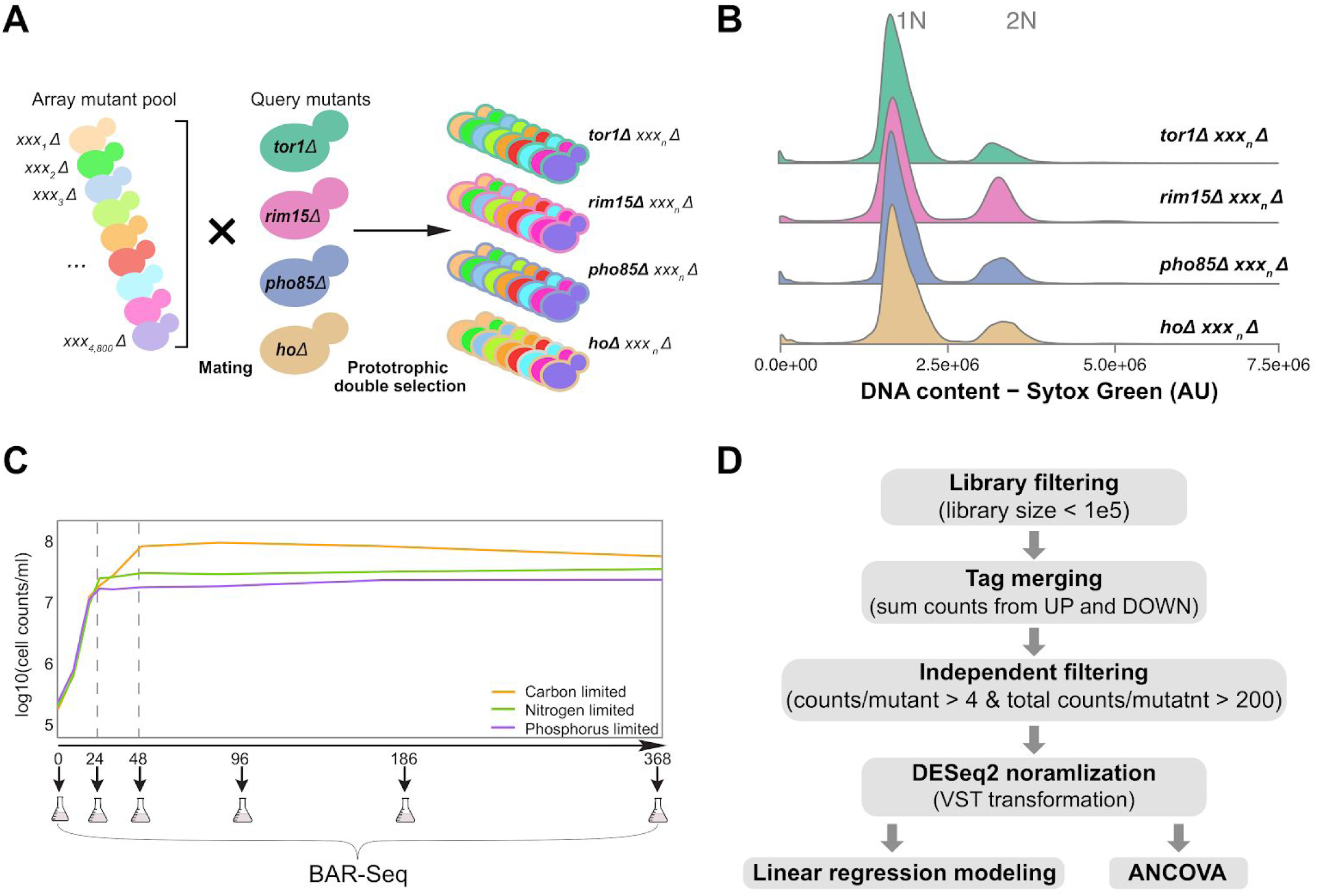
Ploidy confirmation, growth curves and pipeline for data pre-processing. **A)** Ploidy confirmation for double mutant libraries assayed using Syto Green staining. **B)** Bar-seq data pre-processing pipeline for each library/sample. **C)** Growth curve of 12 mutant libraries, 4 different genotypes (HO, RIM15, TOR1, PHO85) × 3 replicates, in different media (orange – carbon restriction, green – nitrogen restriction, purple – phosphorus restriction).

**Figure EV2.**
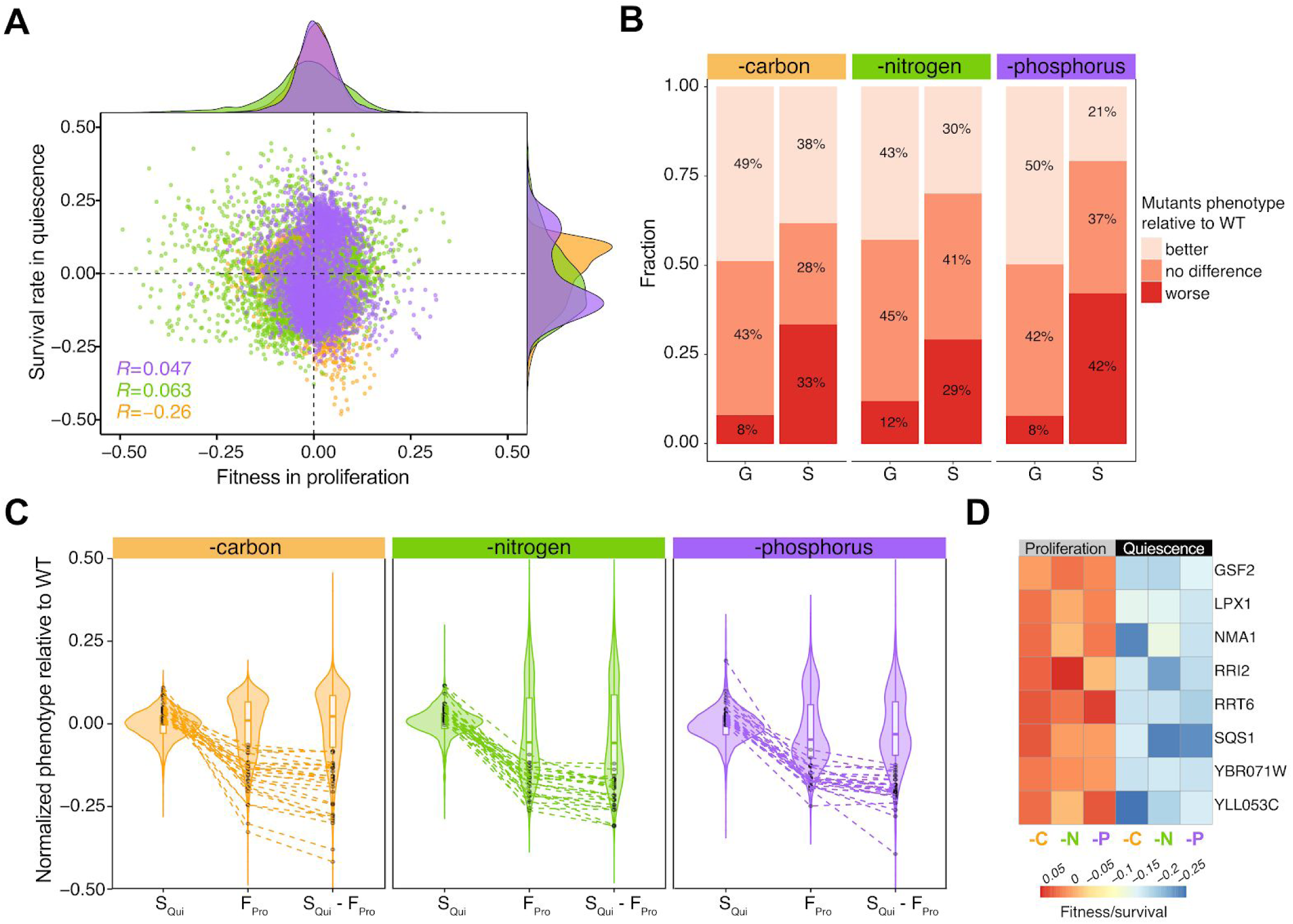
Correlation between fitness in proliferation and survival in quiescence. **A)** Weak correlation between fitness in proliferation and survival in quiescence under three nutrient restricted conditions. The distributions of fitness and survival for thousands of mutant in different conditions are shown on the top (fitness) and right (survival). Pearson correlation score is labeled on the bottom left of plot with p-value < 0.05. All scatter plot, frequency plot and labels are colored based on media types (orange – carbon restriction, green – nitrogen restriction, purple – phosphorus restriction). **B)** The proportion of mutants with different fitness and survival compared to wild type in proliferation and quiescence across three nutrient-restrictions. Proportions were calculated based on the statistics summarized from linear regression modeling. Better than wild type – regression coefficients of those mutants are larger than 0 with corrected p-value less than 0.05; no difference compared to wild type are the mutants with corrected p-value of regression coefficient greater than 0.05; worse than wild type are the mutants whose regression coefficient is less than 0 with corrected p-value less than 0.05. **C)** Criteria used to screen for the core set of genetic factors required for cellular quiescence. Those genes that meet the criteria are connected across distributions of different cellular states (F_Pro_, S_Qui_) and phenotypic difference (S_Qui_ – F_Pro_). These genes share the following features: 1. no significant defects compared to wild type in proliferation, 2. strong survival defects in quiescence, 3. phenotypic difference between proliferation and quiescence is statistical significant. **D)** The 8 genes defined in **Figure 2D** and their corresponding phenotypic readout in different conditions and cellular states.

**Figure EV3.**
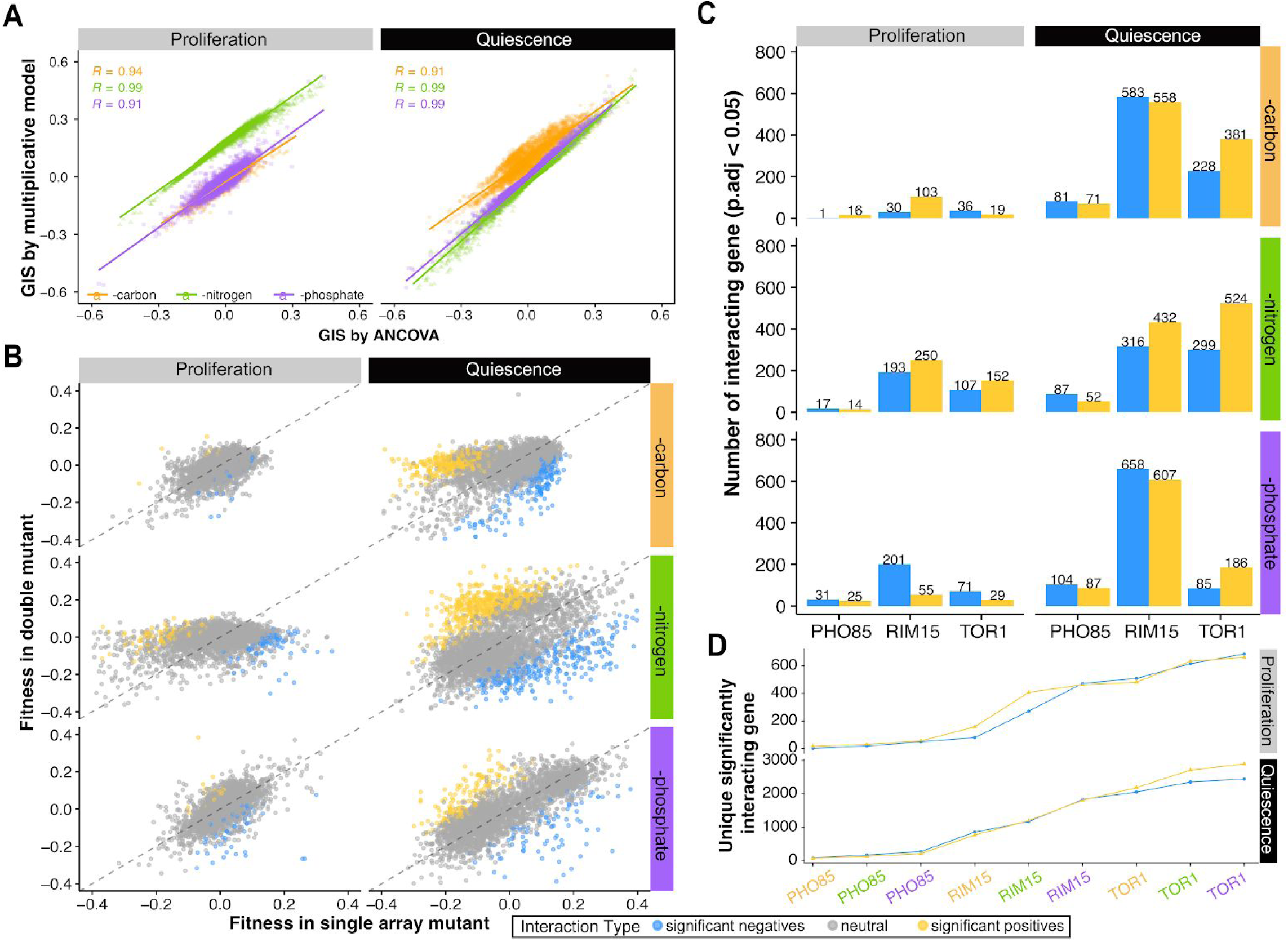
Genetic interaction quantification comparison and summary. **A)** Comparison of genetic interaction score between kinase TOR1 and other non-essential genes estimated by ANCOVA and multiplicative model for different cellular states. Pearson correlation between ANCOVA and multiplicative model calculated GIS is labeled on the bottom left of the plot with p-value < 0.05. A linear regression line is plotted for each condition for each cellular state. Both scatter plot, linear regression line and labels are colored based on media types (orange – carbon restriction, green – nitrogen restriction, purple – phosphorus restriction). **B)** Scatter plot of fitness and survival estimated in double mutation background (tor1Δ0 xxxΔ0: y-axis) and single mutation background (xxxΔ0: x-axis). The dashed diagonal line is colored as grey. A similar trend is found for other query mutants **C)** Quantitative summary of significantly interacting genes (p.val < 0.05) with each kinase in proliferation and quiescence. **D)** Cumulative plot of unique genetic interactions detected with each kinase in three nutrient media (orange – carbon restriction, green – nitrogen restriction, purple – phosphorus restriction).

**Figure EV4.**
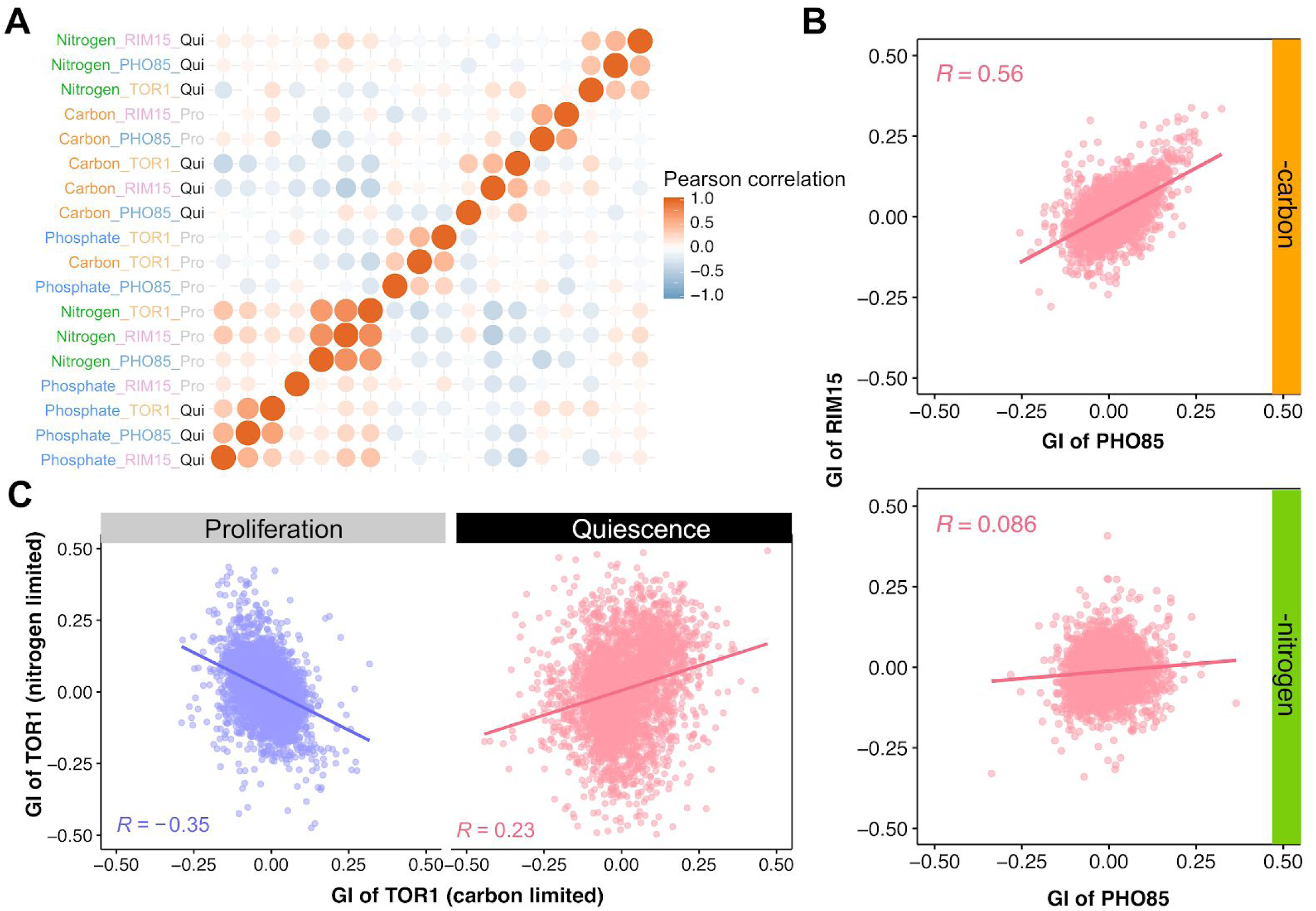
Genetic interaction profiles with kinases across difference conditions. **A)** correlation heatmap of genetic interaction profiles for each kinase under two cellular states in response to different nutritional restriction or starvations. Samples are orders based on hierarchical clustering. **B)** Comparison of genetic interaction profiles between PHO85 and RIM15 in carbon- (top) or nitrogen- (bottom) restricted proliferating cells. Pearson correlation score is labeled in the plot with p-value < 0.05 (color code: pink – positive correlation). **C)** comparison of genetic interaction profiles of TOR1 between different nutrient-restricted and -starved conditions. Calculate pearson correlation is plotted on the bottom right of each panel with p-value < 0.05 (color code: pink – positive correlation, purple – negative correlation).

**Figure EV5.**
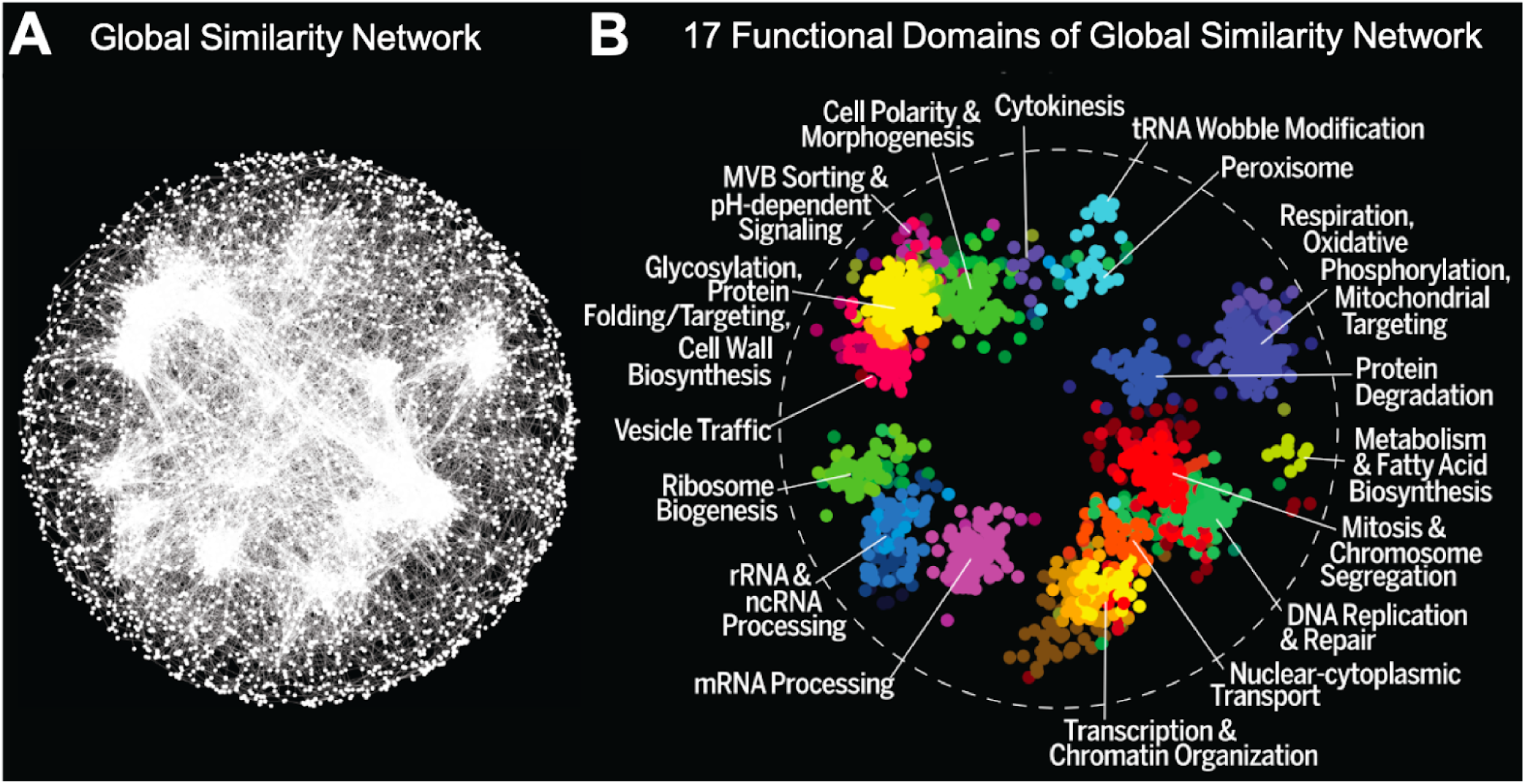
A global network of genetic interaction profiles similarities. **A)** A global genetic profile similarity network encompassing ∼3,971 nonessential and essential genes from Costanzo et al., 2016. **B)** The global similarity network was annotated using the Spatial Analysis of Functional Enrichment (SAFE) (Baryshnikova 2016), identifying network regions enriched for similar GO biological process terms, which are color-coded (Costanzo et al., 2016).

**Figure EV6.**
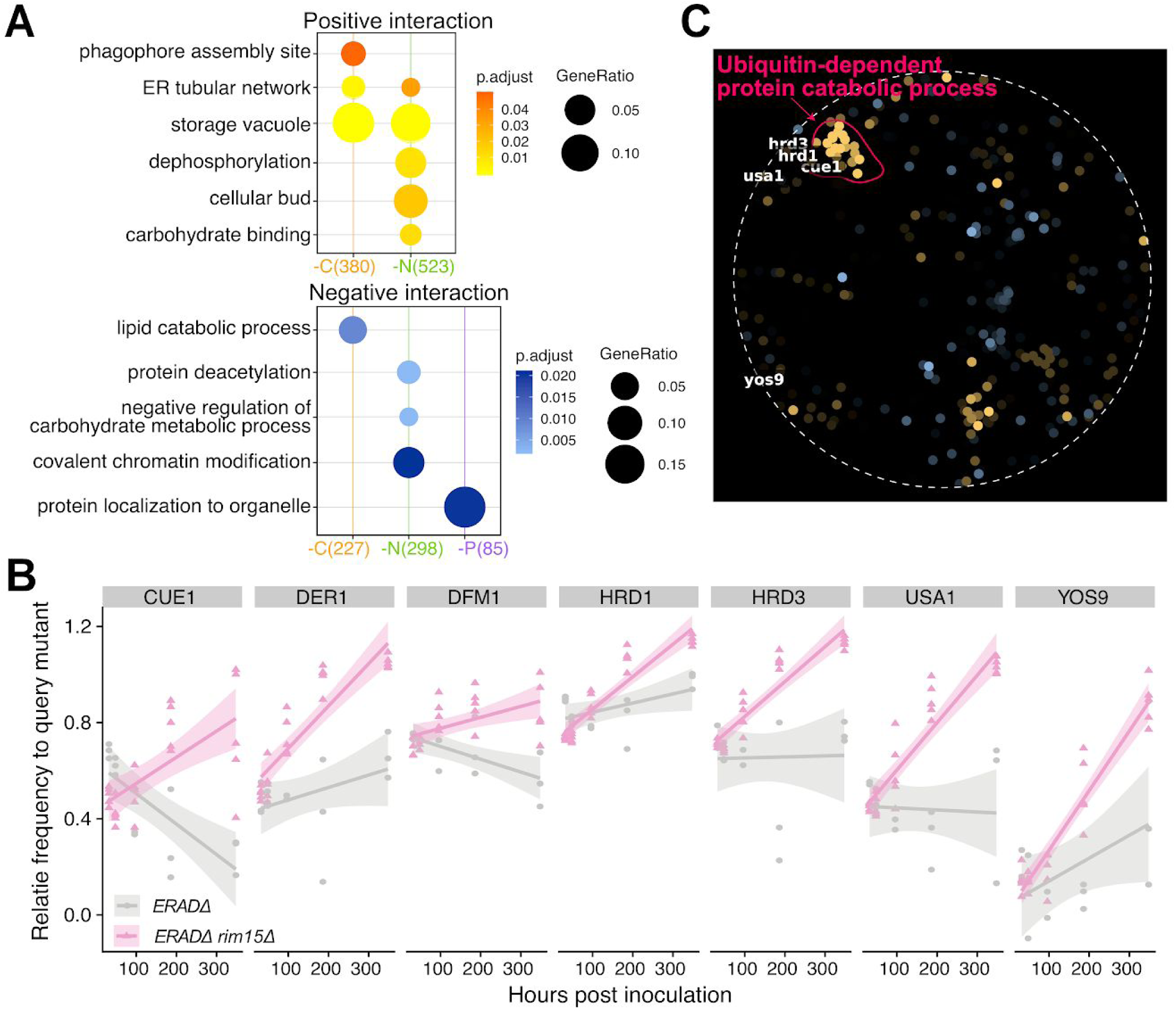
Functional analysis of significantly interacting genes with different kinase. **A)** GOterm enrichment analysis for genes that significantly interact with TOR1 in each different nutrient starvation condition. Only GOterms with significant representation are shown (p.adjust < 0.05). The same color schemes are used to represent different interaction types (yellow – positive, blue – negative). The intensity of the dot color represents the significance, e.g. the lighter the color is, the smaller p-value. The size of the dot represents the gene group size within each term, given the significant interacting genes under each condition (colored parentheses on x-axis). **B)** Relative frequency of each double (*ERADΔ0 rim15Δ0*) and single mutant (*ERADΔ0*) as a function of time in response to nitrogen starvation. **C)** SAFE analysis for genes interacting with RIM15 in nitrogen starvation conditions.

